# Tunneling nanotubes propagate preneoplastic cues in a BMP-dependent manner

**DOI:** 10.1101/2025.11.11.687852

**Authors:** Cristina Cuella Martin, Thanh Thuy Nguyen, Kevin Geistlich, Alexia Bertin, Boris Guyot, Emmanuel Delay, Charline Dalverny, Alice Boissard, Cécile Henry, François Guillonneau, Laurie Tonon, Sylvain Lefort, Eve-Isabelle Pécheur, Véronique Maguer-Satta

**Author notes:** Co-corresponding authors: Eve-Isabelle PÉCHEUR and Véronique MAGUER-SATTA, Centre de Recherche en Cancérologie de Lyon-CRCL, UMR CNRS 5286/Inserm U1052, 28 rue Laënnec, 69373 Lyon Cedex 08, France; 33-4-7878-2907 or 33-4-7878-2806. These authors contributed equally (Thanh Thuy Nguyen, Kevin Geistlich, Eve-Isabelle Pécheur, Véronique Maguer-Satta).

## Abstract

Tunneling nanotubes (TNTs) are thin, actin-rich structures that facilitate long-distance intercellular communication through the transfer of molecules and organelles. While their role in cancer progression is well-established, their involvement in the earliest stages of tumor initiation remains unexplored. Using normal human mammary primary cells, a model of early luminal breast transformation, and mammary organoids, we demonstrate that TNTs become increasingly abundant and elongated during the initial phases of luminal breast cell transformation. Notably, these structures preferentially connect transformed donor cells to non-transformed acceptor cells, establishing an horizontal communication network, favoring the direction of transfer from transformed to non transformed cells. Unbiased proteomics and single-cell RNA sequencing analyses, coupled with (correlative) super-resolution microscopy reveal that, through this network, transformed cells transfer key molecular signals to non-transformed acceptor cells, inducing changes of their proteome and transcriptome landscape within days. Further analyses of acceptor cells reveal dysregulation of the BMP pathway, including increased expression of the Bone Morphogenetic Protein receptor type 1b (BMPR1b), together with functional phenotypic alterations such as the acquisition of anchorage-independent growth, a hallmark of preneoplastic progression. Our findings highlight how TNT-mediated transfer initiates a BMP-dependent cascade of molecular and phenotypic changes in acceptor cells, effectively propagating preneoplastic cues. By elucidating these early events, our findings provide new mechanistic insights into the onset of epithelial transformation in luminal breast cancer, identify the BMP pathway as a key mediator of this process, and advance our understanding of how preneoplastic states emerge and disseminate.

## Introduction

With 2.3 million women affected worldwide and 700,000 deaths per year, breast cancer is the leading cause of cancer-related death in women, with luminal types representing 70% of all cases [1], and ductal carcinoma *in situ* (DCIS) the earliest non-invasive stage. The Bone Morphogenetic Proteins (BMP) pathway is a key actor altered during luminal breast tumorigenesis and further tumor progression [2–5]. Patients with breast cancer exhibit increased expression of BMP ligands and of the BMP receptors BMPR1a and BMPR2, which correlates with a poor prognosis. This indicates that BMP signaling in breast cancer cells remains active during tumor progression and metastasis [2]. In the context of luminal breast tumor, enhanced BMP2 secretion by the niche surrounding mammary stem cells fuels their transformation into cancer stem cells *via* amplification of their response to the kinase BMP receptor type 1b (BMPR1b), without detectable genetic alterations [4,6]. This is concomitant to an interleukin-6 (IL6)-mediated inflammation. This cytokine, among others (IL10, TGFβ) maintains stemness and promotes survival of cancer stem cells, which can drive tumorigenesis and chemoresistance [7–9]. BMPR1b-silenced transformed mammary cells have drastically decreased capacities to initiate tumor formation *in vivo* [4]. However, the early dynamics of BMP-mediated transformation and the evolution of non-transformed mammary cells towards tumor cells remain unexplored.

Tunneling nanotubes (TNTs) are thin, transient actin-based structures enabling long-range intercellular communication [10,11], *via* the transfer of organelles, small molecules, proteins and nucleic acids. TNTs have been implicated in tumor progression, metastasis, and chemoresistance. Cancer cells used these tubes to interconnect over ultra-long distances, disseminate and invade the organ [12]. Also cancer cells linked to neurons by TNTs exhibited enhanced metastatic capacities [13–15]. Yet the role of TNTs at earliest stages of tumorigenesis is unknown.

While BMPs and TNTs have independently been associated with breast cancer, their potential interaction is unexplored. Furthemore, the nature, directionality, and functional consequences of cell/cell communications at the tumor initiation stage remain poorly understood.

In this work, we aimed to delineate the interplay between the BMP pathway and TNT-mediated communication, to shed light on the earliest steps of transformation of (immature) mammary cells.

## Results

### TNT formation is increased during mammary epithelial stem cells transformation

Early BMP-driven transformation dynamics was explored by using our clinically-relevant MCF10A-derived stem cell models, obtained by chronic exposure to IL6 and BMP2 (MC26 and M1B26; Figure 1A) [4,6]. Sorting of BMPR1b-expressing cells prior to cytokine treatment, followed by soft agar clones selection, led to a quicker establishment of transformed cells (M1B26), with greater *in vitro* anchorage-independent growth and *in vivo* engraftment than in non BMPR1b-sorted cells (MC26) [4]. These models display DCIS molecular features and robustly recapitulate the *in vivo* BMP pathway alterations [4,6]. Super-resolution Airyscan confocal microscopy observations of non-transformed control (MCF10A-CT) and transformed cells (MC26, M1B26) revealed long-range contacts *via* long thin F-actin-rich TNTs [16] (Figure 1B). Although TNT quantification requires low-confluence culture conditions [17], long-range TNTs were also detectable at high cell density (Supplementary Figure S1A). While the average length of TNTs in non-transformed or transformed cells did not significantly vary (Figure 1C left), M1B26 transformed cells exhibited over 4-fold more TNTs as non-transformed cells (Figure 1C middle), and the proportion of long TNTs increased ∼fivefold with transformation (Figure 1C right). This apparent discrepancy, between a high proportion of long TNTs and a similar average TNT length upon transformation, could be linked to the abundance of shorter TNTs when all TNTs are considered together. Importantly, transformed cells could establish TNT connections with non-transformed primary epithelial mammary cells (Figure 1D), isolated from healthy human breast samples. On the contrary, neither TNT number nor length varied in non-transformed cells exposed to conditioned medium of M1B26 cells (Figure 1C left, Supplementary Figure S1B-C), suggesting that secreted factors are not sufficient to promote TNTs in non-transformed cells.

**Figure 1.**
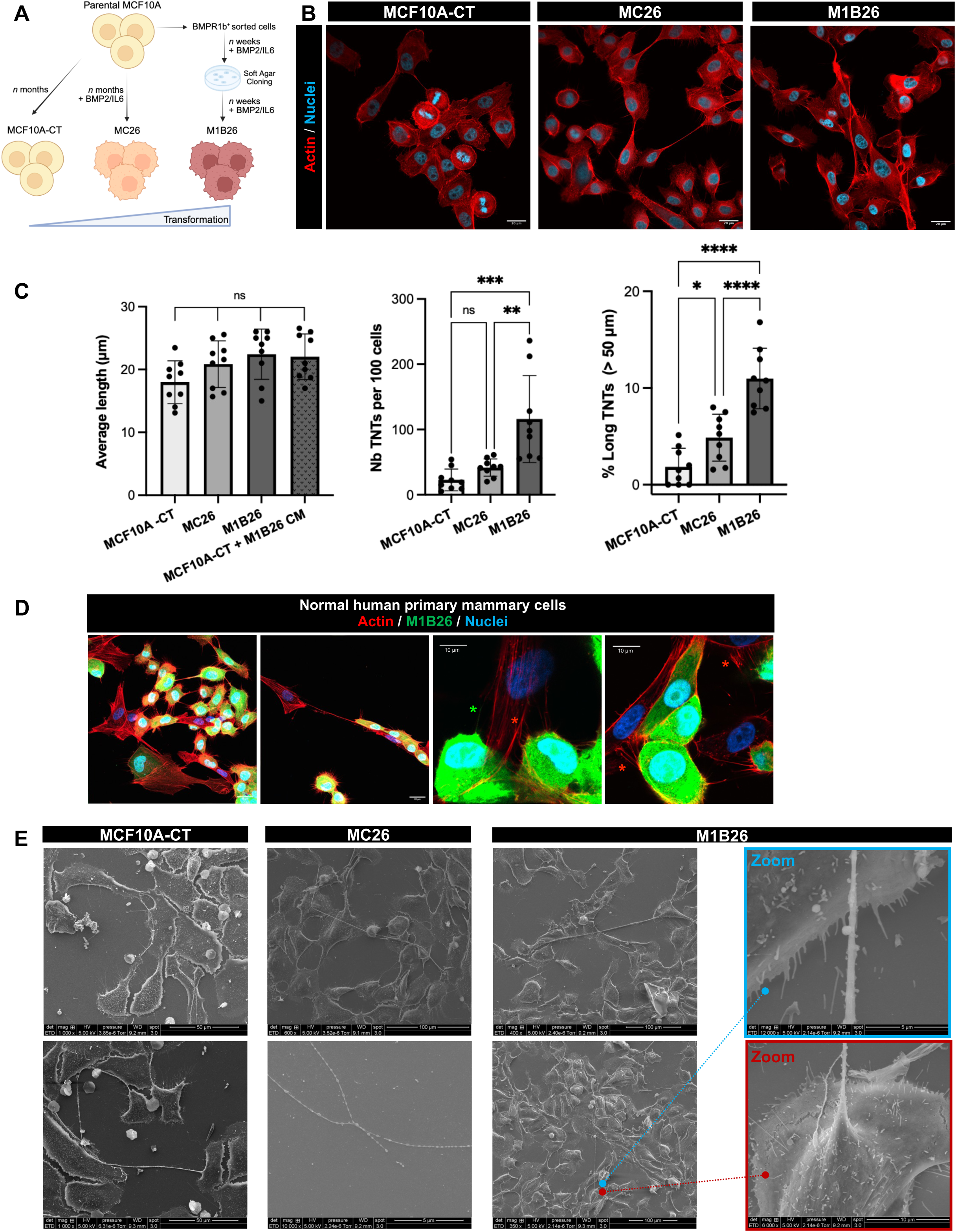
TNT formation is enhanced upon mammary cell transformation. A, Schematic of our MCF10A-derived cell lines [6]. B, Immunofluorescence images of actin (red) in TNTs, observed by super-resolution Airyscan confocal microscopy with an 40x oil objective. Nuclei, Hoechst labeling (blue). Scale bars, 20 μm. C, From left to right: quantification of TNT average length for indicated cell lines and conditions (MCF10A-CT+M1B26 CM : MCF10A-CT cells incubated for 48h with conditioned medium of M1B26 cells); TNT number (> 8 μm) quantification *per* 100 cells; % of long TNTs (> 50 μm); n = 9 biological replicates for each condition. Ten Z-stacks of 10 sections each were taken *per* condition, and 100 cells minimum *per* experiment were counted. TNTs were counted and measured in each Z-stack. Error bars depict SD. P-values were calculated with one-way ANOVA with post-hoc Tukey’s Honest Significance Difference test (ns, not significant; * p < 0.05; ** p < 0.01; *** p < 0.001; **** p < 0.0001). D, Images of TNTs linking M1B26 (GFP, green) to normal human primary mammary cells. Actin, red; nuclei, blue. Asterisks: emerging from M1B26 (green); between both cell types (red). Scale bars: left, 20 μm; right, 10 μm. E, Scanning electron microscopy images of TNTs of MCF10A-CT, MC26 and M1B26 cells. Scale bars MCF10A-CT: 50 μm; MC26: 100 and 5 μm; M1B26: 100 μm in low mag, 10 or 5 μm in zooms. Colored lines indicate regions where zoom was performed.

Scanning electron microscopy revealed that TNTs in transformed cells were often branched, with ramifications, multiple connections and bulges (Figure 1E, Supplementary Figure S1D) [10,16]. TNTs did not adhere to the substrate (Figure 1E zooms, Figure 2A, Movie S1), although it cannot be excluded that some tiny “threads”, spreading out from TNTs, might play a role in adherence (Supplementary Figure S1D M1B26 cells). TNTs were also observed within 3D structures such as acini, the basic functional units of breast glandular tissue (Figure 2B-C, Movie S2). While some TNTs were open-ended (Figure 2D, Supplementary Figure S2A), allowing a cell-cell cytoplasmic continuum, some close-ended TNTs featuring a Gap junction at one tip were seen (Supplementary Figure S2B). Moreover, TNTs formed thin and elongated membrane structures, with an average diameter of 220 ± 23 nm. They also contained mitochondria [18] (Figure 2D-E, Movie S3), and microtubules [10,19], as illustrated by tubulin staining (Supplementary Figure S2C).

**Figure 2.**
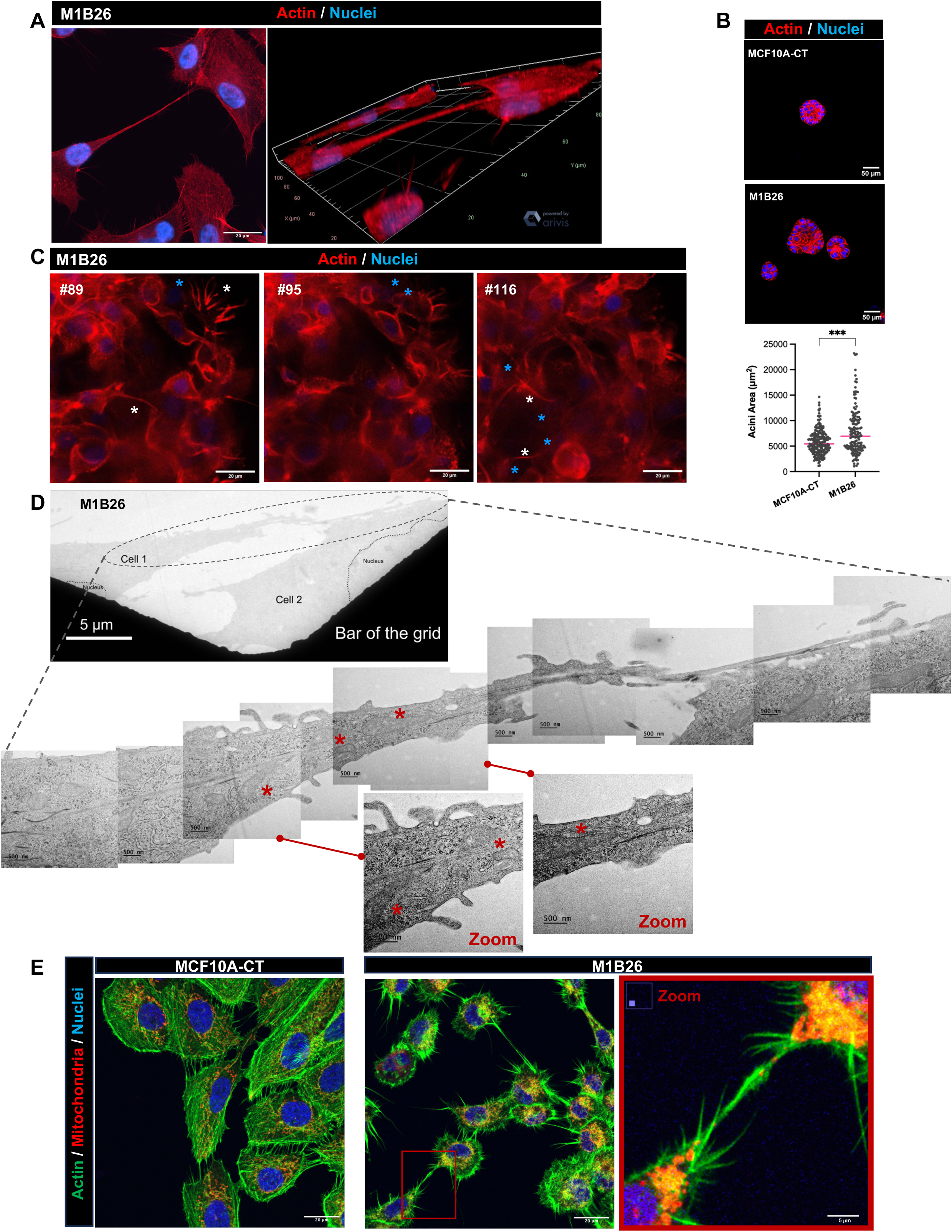
TNTs are non-adherent membrane structures, exist in 3D and contain mitochondria. A, Left, immunofluorescence image of actin (red) in a TNT linking M1B26 transformed cells (scale bar 20 μm). This image was extracted from a Z-stack of 20 images, taken by super-resolution Airyscan confocal microscopy with an 40x oil objective; right, 3D-projection of this stack, obtained with Zeiss ZENlite3.6 software (see Movie S1). Nuclei, blue. B, Immunofluorescence images of actin (red) in mammary acini grown from MCF10A-CT or M1B26 cells (scale bars, 50 μm). Bottom, acini area (in µm^2^). Statistiscal significance was calculated using a one-way ANOVA with post-hoc Tukey test (*** p < 0.001). C, Selected snapshot numbers (#) from Movie S2, with actin and nuclei labeling (red and blue, respectively). White and blue asterisks, TNTs linking cells and nuclei of corresponding cells, respectively. D, Transmission electron microscopy images of an open-ended TNT linking two M1B26 cells, with mitochondria (red asterisks). Scale bars, 5 μm (low mag), 500 nm (high mag). Two images were zoomed (linked with a red line to original image) and their contrast enhanced, to better visualize mitochondria within TNT (red asterisks). E, Immunofluorescence images of mitochondria (Mitotracker^®^-red) in MCF10A-CT and M1B26 cells. Actin, green; nuclei, blue. Scale bars, 20 μm, 5 μm in zoom.

This indicates that early mammary cell transformation leads to an enhanced ability of transformed cells to connect to their homo- or hetero-typic neighbors through TNTs.

### Horizontal transfer of cellular material *via* TNTs from transformed to non-transformed cells

To investigate how TNTs connect cells, we co-cultured mCherry-expressing MCF10A-CT with GFP-expressing M1B26 cells, seeded at low density [17]. Both M1B26 transformed (Tf) and MCF10A-CT non-transformed (NoTf) cells formed TNT structures, connecting Tf to Tf, NoTf to NoTf and Tf to NoTf cells (Figure 3A). Quantification of TNTs contacts revealed that connections were more frequent and abundant from Tf towards NoTf cells than in the opposite direction (Figure 3B,C with schematic). Fluorescent material from GFP-tagged Tf cells accumulated within the cytoplasm of NoTf cells (Figure 3D), suggesting a directional transfer *via* TNTs, Tf cells as donors and NoTf cells as acceptors (Figure 3E; Supplementary Movie S4; Supplementary Figure S3A). To assess the proportion of cellular exchange between cells, mCherry-NoTf cells were cocultured with GFP-Tf cells for 8 days. Flow cytometry analysis revealed a higher proportion of mCherry-NoTf cells exhibiting green fluorescence uptake compared to GFP-Tf cells acquiring red fluorescence (Supplementary Figure S3B), indicating a preferential Tf-to-NoTf horizontal transfer (Figure 3F-H). Switching fluorophores between Tf and NoTf cells yielded similar results (Supplementary Figure S3C,D), and this transfer increased over time (Supplementary Figure S3E-I). To evaluate the contribution of soluble factors and extracellular vesicles to cell-cell communication, normal human primary mammary cells and Tf cells were cultured either separated by a physical barrier (Transwell^™^), which prevent TNT formation, or in the presence of Tf cells conditioned medium (CM, Figure 3I). Transwell^™^ separation reduced the number of acceptor cells in both transfer directions (Figure 3J,K; Supplementary Figure S3J for MCF10A cells), whereas conditioned medium alone did not induce detectable fluorescence exchange (Supplementary Figure S3K,L for normal primary cells). Second, we performed cocultures of NoTf with Tf cells, in the presence or absence of either TNTi, a previously described inhibitor of TNT formation [20,21], or of cytochalasin D to inhibit actin polymerisation. Both inhibitors, without affecting cellular viability (Supplementary Figure S5A), drastically decreased TNT number (Supplementary Figures S4, S5B-C), while inhibiting the transfer of mitochondria between mitotracker-stained Tf cells and NoTf cells (Supplementary Figure S4C, Movie S5). Third, we blocked exosome biogenesis in cocultures of Tf and NoTf cells, by using GW4869 [22]. In agreement with literature, Tf cells secreted higher amounts of EVs than NoTf cells, as evidenced by higher amounts of CD63 and CD9 in EVs-enriched supernatants from Tf cells (Supplementary Figure S5D-F). Inhibiting exosome biogenesis with GW4869, in cocultures of fluorescently-labeled NoTf and Tf cells, consistently led to decreased levels of CD63 and CD9 in the cell supernatant (Supplementary Figure S5G, I-J), without affecting cell viability (Supplementary Figure S5H). In contrast, as revealed by flow cytometry, the uptake of fluorescent material, in proportion and intensity, remained higher in the Tf-to-NoTf direction than in the other direction, whether or not GW4869 was present (Supplementary Figure S5K-L).

**Figure 3.**
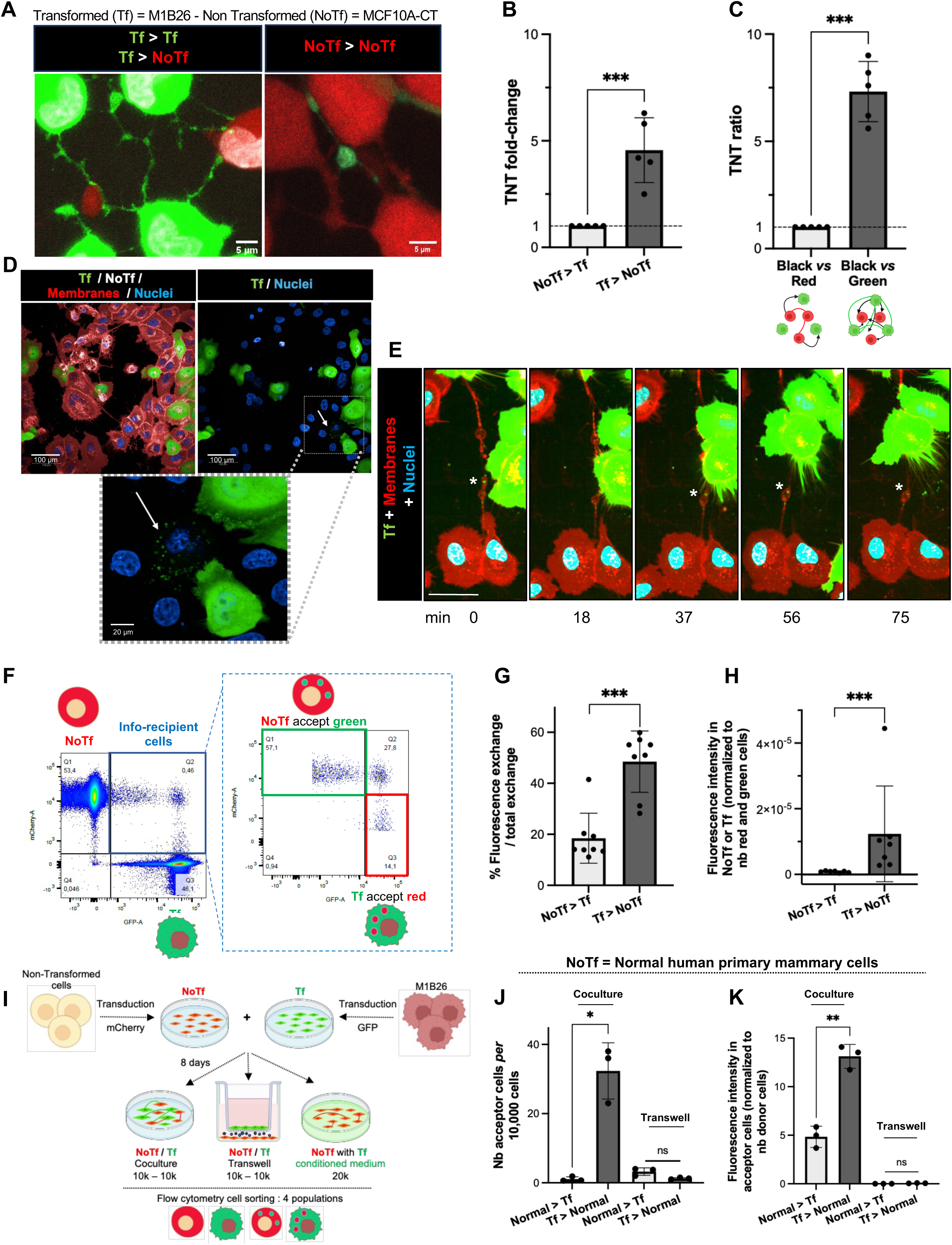
TNTs link transformed and non-transformed mammary cells, favoring an horizontal transfer of information. A, Fluorescence images of GFP-labeled-M1B26 (transformed - Tf; green) and mCherry-labeled MCF10A (non-transformed - NoTf; red) cells, after 48 h in coculture. Scale bars, 5 μm. B,C, Relative number of TNTs linking NoTf to Tf, or Tf to NoTf cells (B) and TNT ratio defined as 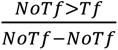 (black arrows *vs* red lines **in schematic**) or 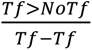 (black arrows *vs* green lines **in schematic**) after 48h in coculture (C). Z stacks of 10 sections each were taken *per* condition and 100 cells minimum were counted. TNTs were counted and measured in each Z-stack. Error bars depict SD. Statistical significance was calculated using an unpaired Student’s two-sided Welch’s t test (n = 5 biological replicates; *** p < 0.001). D, Fluorescence images of material transfer (green dots) from Tf (green) to NoTf cells. Membranes, red; nuclei, blue. Scale bars: 100 μm, in zoom 20 μm. E, Time-lapse snapshots of green material trafficking within a TNT bulge (*), from a (green) Tf to NoTf cells after 48 h in coculture. Membranes, red; nuclei, blue. Scale bar, 100 μm; images recorded for 75 min, with one image every 18 min (see Supplementary Movie S4). F, Flow cytometry analysis of an 8-day coculture of green Tf and red NoTf cells, with a schematic of NoTf accepting material from Tf cells (top: NoTf accept green, red cell with green dots), and Tf cells accepting material from NoTf cells (bottom: Tf accept red, green cell with red dots). G,H, Quantification of the percentage of exchange (G; n = 8 biological replicates) and fluorescence intensity (H; n = 7 biological replicates), for indicated transfer. Error bars depict SD. Statistical significance was calculated using an unpaired Student’s two-sided Welch’s t test (*** p < 0.001). I, Schematic of the experimental design of coculture experiments. J,K, Quantification of the number of acceptor cells *per* 10,000 cells (J), and fluorescence intensity (K), in standard or Transwell™ cocultures, for indicated transfer involving normal human primary mammary (normal) and M1B26 cells (Tf). Error bars depict SD. Statistical significance was calculated using an unpaired Student’s two-sided Welch’s t test (n = 3 biological replicates; ns, not significant; * p < 0.05; ** p < 0.01).

Taken together, these findings support that horizontal material transfer preferentially occurs *via* TNTs from Tf toward NoTf cells, rather than through secreted factors or extracellular vesicle-mediated communication.

### TNT-mediated transfer modulates proteomic features of acceptor cells

To investigate the molecular consequences of intercellular material transfer, we performed a proteomic analysis of distinct cell populations isolated from long-term cocultures. Fluorescently-labeled cells cultured for 8 days (Figure 4A), were sorted by flow cytometry into four populations: non-transformed (NoTf), transformed (Tf), non-transformed cells that received fluorescent material (NoTf accept), and non-transformed cells from the same coculture that did not receive any material (NoTf no accept). Unbiased proteomic screening revealed that NoTf accept cells displayed a distinct proteomic signature compared to NoTf no-accept cells from the same coculture and NoTf cells in monoculture (Figure 4B, Supplementary Figure S6A). Gene set enrichment analysis further showed increase of pathways related to mitochondrion organization and activity, or transport and localization of intracellular organelles, macromolecules and proteins (Figure 4C, Supplementary Figure S6B). Of note, the most abundant proteins in NoTf accept cells were involved in intracellular trafficking and lipid metabolism (RAB11B, MEST, PIGA), cell growth and morphology maintenance (TRIQK), membrane protrusion formation (ATP8B1), actin filament bundling (LCP1 / Plastin-2), and the import of NAD^+^ into mitochondria (SLC25A51, SLC25A52) (Supplementary Table S1). Four proteins, Rab-11B, Plastin-2, MEST and TRIQK, were markedly more abundant in NoTf accept cells compared to NoTf no-accept cells from same cocultures (Figure 4D). Preventing TNT formation using Transwell inserts abolished the molecular changes observed in NoTf accept cells, which instead share common features with to NoTf in monoculture (Supplementary Table S2, Supplementary Figure S6C). By contrast, NoTf accept cells from direct cocultures exhibited a higher abundance of early inflammatory markers (SERPINA3, CD14), and extracellular matrix-related genes (LAMC2, CHI3L1, LAMB3, IGFBP6) (Supplementary Table S2, Supplementary Figure S6D). Interestingly, NoTf accept cells from direct cocultures also displayed high amounts of MACROD1, which plays a role in estrogen signaling and whose overexpression is correlated to enhanced cellular invasive capacities. These findings indicate that direct contacts with transformed cells induce profound changes in the proteomic landscape of non-transformed acceptor cells, similar to those observed in early transformed cells (M1B26 cells).

**Figure 4.**
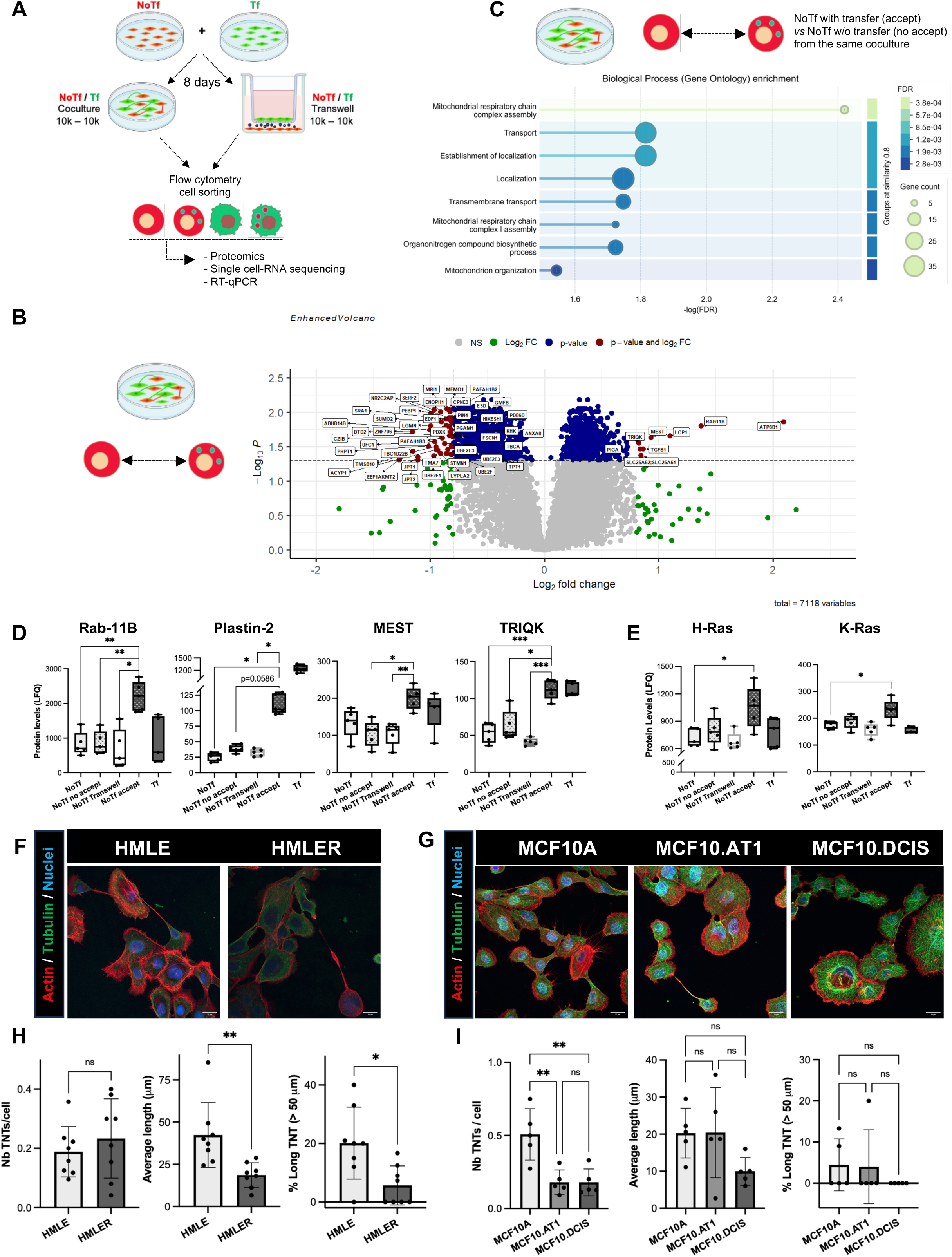
**TNT-dependent phenotypic changes of non-transformed cells acceptor of material from transformed cells**. A, Schematic of the experimental design: 4 cell populations, sorted by flow cytometry (NoTf, Tf, NoTf in coculture without fluorescent transfer [non-transformed non-acceptor cells -NoTf no accept], NoTf in coculture with fluorescent transfer [non-transformed acceptor cells - NoTf accept]), underwent proteomic analyses and RT-qPCR. B,C, Volcano plot (B) showing Log_2_ enrichment of individual proteins in NoTf accept compared to NoTf no accept cells from the same coculture (see schematic). Adjusted p-value (with Benjamini-Hochberg procedure) threshold was set at 0.05 and 0.8 for Log_2_ fold change. C, To obtain a larger pool of hits, it was lowered to 0.5, generating a list of 193 proteins upregulated in NoTf accept cells, used to generate the GOBP subset with STRING (https://string-db.org; RRID:SCR_005223). D,E, Box and whiskers plots of Log_2_LFQ values (Protein Levels) of Rab-11B (RAB11B), Plastin-2 (LCP1), MEST (MEST), TRIQK (TRIQK) (D), and H-Ras and K-Ras (E), for indicated sorted cell populations. Statistical significance was calculated using one-way ANOVA with Dunnett’s multiple comparison test (n = 5 biological replicates; * p < 0.05 ; ** p < 0.01 ; *** p < 0.001 ; **** p < 0.0001). For clarity, comparisons with Tf cells were omitted, and in (E), only significant comparison are shown. F,G, Immunofluorescence images of actin (red) and tubulin (green) in (F) HMLE and HMLER cells, and (G) MCF10A, MCF10.AT1 and MCF10.DCIS cells. Nuclei, blue. Scale bars = 20 μm. H,I, From left to right: quantification of TNT number *per* cell, average length, and percentage of long TNTs in indicated cell lines. Ten Z-stacks of 10 sections each were taken *per* condition and 100 cells minimum *per* experiment were counted. TNTs were counted and measured in each Z-stack. Statistical significance was calculated using an unpaired Student’s two-sided Welch’s t test (n = 8 and n = 5 biological replicates for HMEC-derived and MCF10A-derived cell lines, respectively; ns, not significant; ** p < 0.01 ).

To determine whether TNT formation is similarly affected in other independent models of mammary epithelial cell transformation, we next used either human mammary epithelial cells immortalized with SV40-L antigen and hTERT, subsequently transformed with H-Ras [23] (Figure 4E-F), or H-Ras-transformed immature MCF10A-derived AT1 and DCIS cells (Figure 4G-H). Unlike M1B26 cells, neither model displayed an increased number of TNTs or greater proportion of long TNTs (Figure 4E-H), suggesting that Ras-driven transformation does not promote TNT formation and may rely on mechanisms distinct from the combined inflammation- and BMP-dependent processes implicated in luminal breast cancer clinical occurrence [4,24].

### TNT-mediated transfer impacts the BMP pathway of acceptor cells, while BMPR1b transits in TNTs

Since (transmembrane) transport was one of the most impacted processes (Figure 4B), and given that the BMP pathway is involved in luminal transformation *in vivo* [4,5], we examined whether this pathway was affected in NoTf acceptor cells. We focused on the canonical SMAD cascade, and among non-canonical MAPK downstream cascades, involved in BMP signaling among other pathways, on p38 (MAPK14), because the p-p38/p38 ratio exhibited the highest level of expression in Tf cells (Supplementary Figure S7A-B). This choice is further supported by the established role of p38 MAPK as a key regulator of oncogenic stress responses and transformation in mammary epithelial cells [25]. A limited set of six proteins involved in these cascades, among others, was extracted from our proteomic screening (Supplementary Table S3), and assessed for their expression across sorted cell populations post-TNT connections (Figure 4A). Several proteins involved in the BMP pathway, including ID1, Nup214 [26], Spartin, and USP15 [27], were consistently up- or down-regulated in NoTf acceptor cells, reaching levels comparable to those observed in Tf cells (M1B26), exclusively under conditions in which TNT could form (Figure 5A). ATF2, a transcription regulator positioned at the BMP pathway/p38 cascade interface, was under-expressed in NoTf acceptor cells, at levels comparable to those in Tf cells. Conversely Rac1, part of a complex involved in actin bundling and stabilization, regulated by p38, was overexpressed in NoTf cells that received material *via* TNTs (Figure 5A). At the level of BMP signaling elements, the BMPR1b receptor was localized within TNTs, particularly in bulges, in normal human primary and model mammary cells (Figure 5B). Correlative light (scanning) electron microscopy (CLEM) enables to distinguish intra-from extra-cellular epitopes by overlaying optical and scanning electron images of the same region. Airyscan confocal imaging revealed a continuum of BMPR1b in cell bodies, as well as in or on TNTs. However, electron microscopy did not reveal any extracellular BMPR1b epitopes at TNTs (Figure 5C). This suggests an extracellular localization in the cell body and an intraluminal presence in TNTs, consistent with BMPR1b trafficking through/within TNTs (see below Figure 6A).

**Figure 5.**
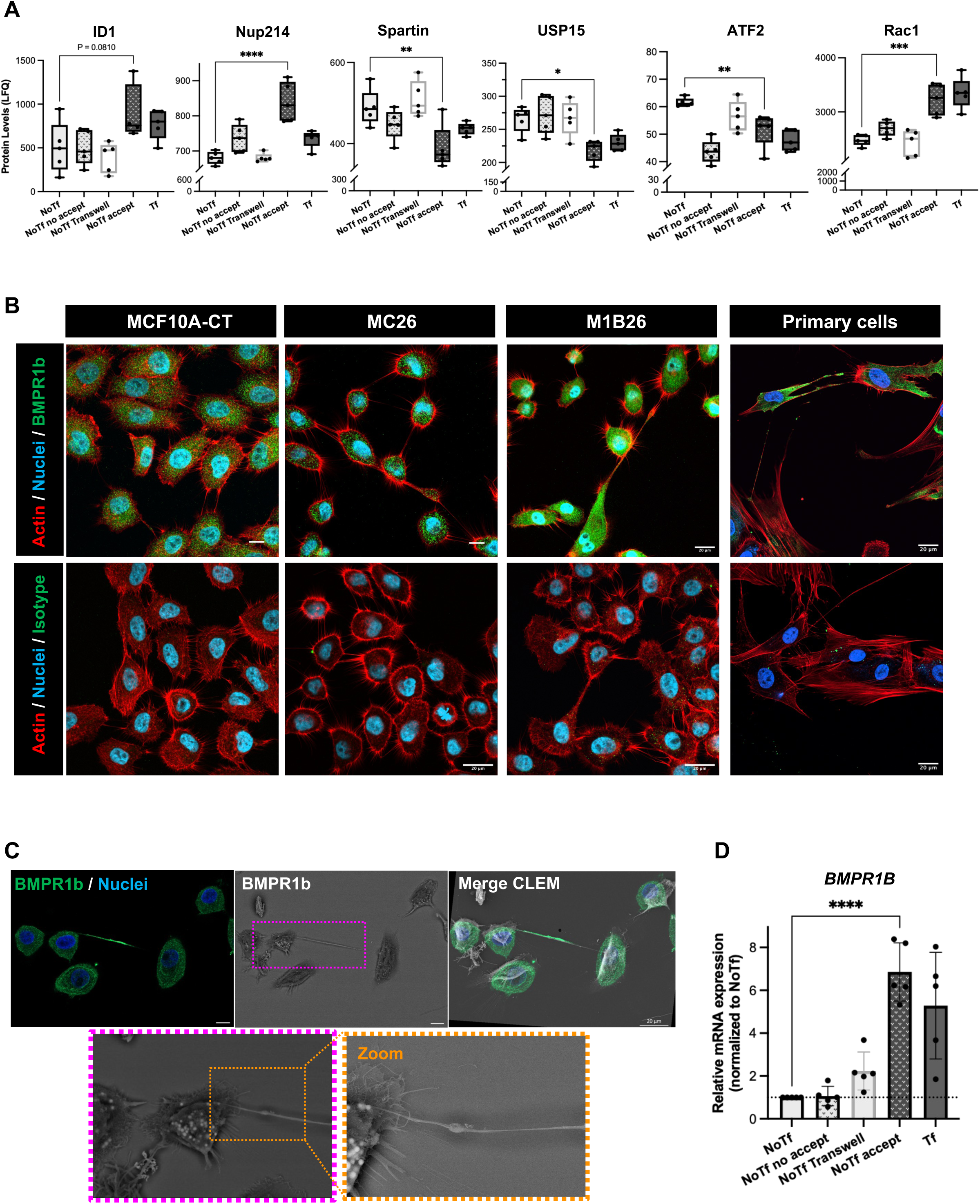
Expression of actors of the BMP signaling pathway is enhanced in acceptor cells, and BMPR1b is present in TNTs. A, Box and whiskers plots of Log_2_LFQ values (Protein Levels) of ID1, Nup214, Spartin, USP15, ATF2 and Rac1, for indicated sorted cell populations. Statistical significance was calculated using one-way ANOVA with Dunnett’s multiple comparison test (n = 5 biological replicates; * p < 0.05; ** p < 0.01; *** p < 0.001; **** p < 0.0001). For clarity, only comparison between NoTf and NoTf accept cells is shown. B, Immunofluorescence images of actin (red), BMPR1b (green) or corresponding isotype (green) in indicated cells. Nuclei (blue). Scale bars, 20 μm. C, Correlative light scanning electron microscopy (SEM) of BMPR1b, immunostained with Alexa Fluor^®^488-Fluoronanogold^™^ (green) in M1B26 cells. From left to right: Airyscan confocal, scanning electron microscopy, zooms, correlative (overlay). Scale bars, 10 μm, unless otherwise indicated. D, RT-qPCR measurement of BMPR1b mRNA in indicated cell populations at day 8, the expression in NoTf cells (monocultures) set at 1. Statistical significance was calculated using one-way ANOVA with post-hoc Tukey’s Honest Significance Difference test (n = 5 biological replicates). For clarity, only comparison between NoTf accept and NoTf cells is shown (**** p < 0.0001).

**Figure 6.**
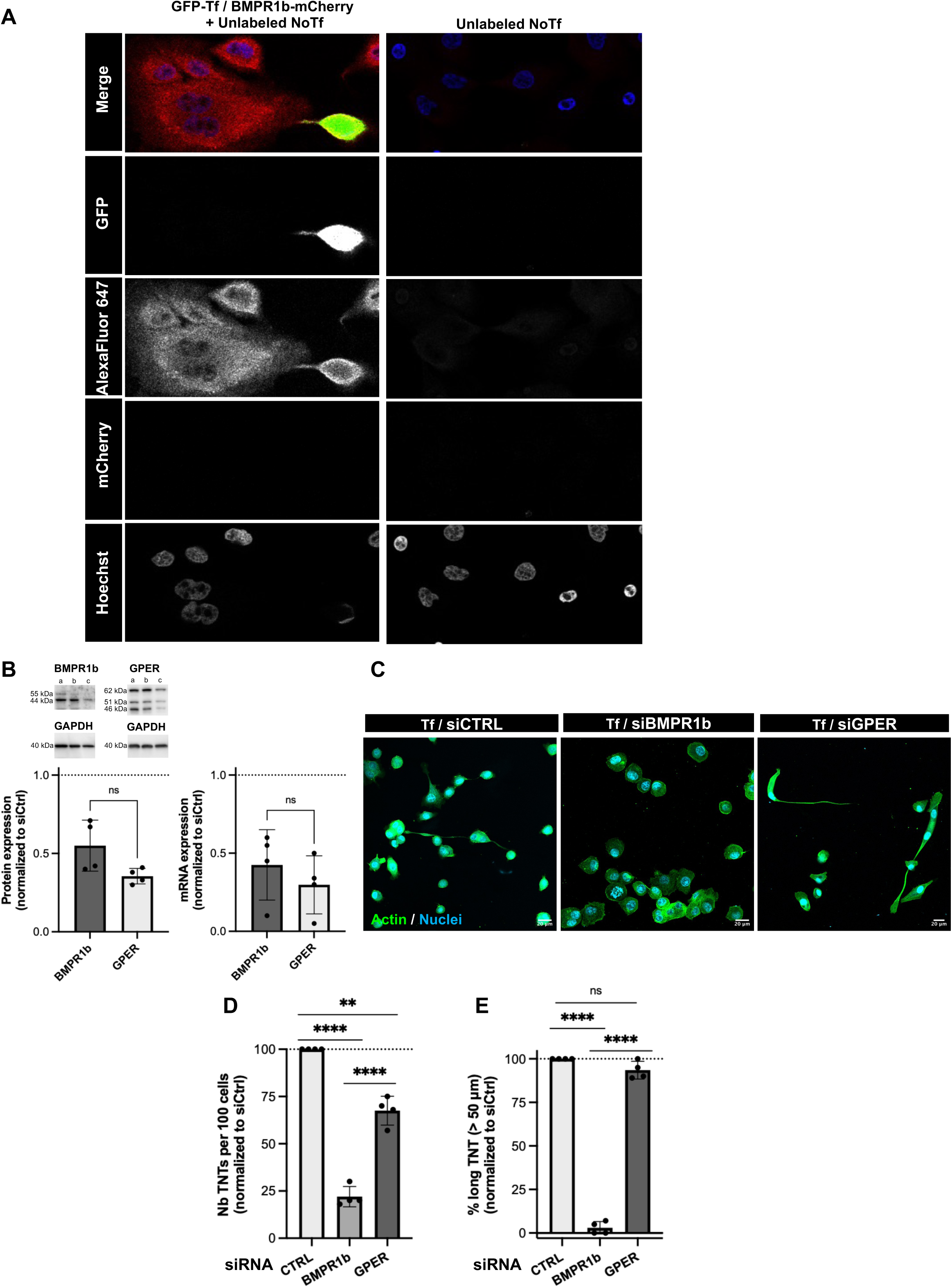
BMPR1b transits within TNTs and contributes to their formation. A, left panels: Immunofluorescence of mCherry-BMPR1b, immunostained with AlexaFluor^®^647 secondary antibody (red), expressed by GFP-labeled Tf cells (green) grown for 48h with NoTf cells (unlabeled); right panels: similar experiment on unlabeled NoTf cells alone. Nuclei, Hoechst labeling (blue). Imaging was performed on a confocal Leica Stellaris 5 microscope, with a 40x oil objective. All fluorescent channels were split for better visualization. B, top panel, Representative immunoblots of BMPR1b and GPER in Tf cell lysates : a, untreated cells; b, cells treated with control siRNA; c, cells treated with targeting siRNA. Bottom panels, BMPR1B and GPER transcript and protein expression upon their silencing, with siCTRL set at 1 (n = 4). C, Immunofluorescence of actin (green), after transfection with siRNA pools targeting *BMPR1b* (siBMPR1b) or GPER (siGPER). Non-targeting siRNA: siCTRL. D-E, TNT number (> 8 μm) *per* cell (D) and % of long TNTs (> 50 μm) (E) upon indicated silencing, normalized against siCTRL set at 100%. Error bars depict SD. Statistical significance was calculated using one-way ANOVA with post-hoc Tukey’s Honest Significance Difference test (n = 4 biological replicates; ns, not significant; ** p < 0.01; **** p < 0.0001).

The four sorted cell populations (Figure 4A) were then analyzed for BMPR1b mRNA expression. BMPR1B levels were approximately 8-fold higher in NoTf accept cells previously connected *via* TNTs, compared to an only ∼2-fold increase when TNT formation was impeded through Transwell assay (Figure 5D).

Whether BMPR1b would transit within TNTs was investigated by generating GFP-labeled Tf cells expressing an mCherry-tagged version of BMPR1b, further co-cultured with unlabeled NoTf cells. After 48h of coculture, mCherry-labeled BMPR1b signal was detected in the cytoplasm of NoTf cells (Figure 6A, Supplementary Figure S7C). In contrast, when cocultures were treated with cytochalasin D prior fixation, inhibiting TNT formation and elongation (see Supplementary Figures S4, S5A-C), no cytoplasmic mCherry signal was detected in NoTf cells, indicating of a blockade of BMPR1b transfer (Supplementary Figure S7C). These observations support the possibility that BMPR1b could transit in TNT-like structures, from Tf to NoTf cells.

We next assessed whether BMPR1b would play a role in TNT formation, using G protein-coupled estrogen receptor 1 (GPER), which has not been implicated in TNT formation/function, as a negative control. Although BMPR1B silencing was only partial (Figure 6B), it markedly reduced TNT number in Tf cells (Figure 6C-D), and drastically reduced the formation of long TNTs (Figure 6E). In contrast, GPER knockdown, although as effective as BMPR1B silencing, had limited effect on TNT number and no significant effect on TNT length (Figure 6C-E). Importantly, BMPR1B silencing also impaired both migration and invasion capacities of Tf cells, without affecting their viability (Supplementary Figure S8A-E). Together, these findings support a role for BMPR1b in TNT formation and elongation, particularly in the generation of long TNTs.

Next, the potential contribution of BMPR1b to TNT-mediated transfer of material between Tf and NoTf cells was investigated by silencing BMPR1b solely in the GFP-labeled Tf cell population (see Materials & Methods, and Supplementary Figure 8F). After their coculture with unlabeled NoTf cells, Tf cells appeared round and devoid of TNTs (Supplementary Figure 8G green asterisk, Movie S6), as already observed (Figure 6C). Importantly, these cells proved unable to transfer mitochondria to NoTf cells. In contrast, mitochondria transfer *via* TNT freely occurred between NoTf cells in the same coculture, not silenced for BMPR1b (Supplementary Figure 8G white asterisks, Movie S6). This further supports the notion that BMPR1b would be involved in TNT formation, and that it would contribute to TNT-mediated transfer of material from Tf to NoTf cells. Together, these findings: (i) indicate that TNT–mediated material transfer from Tf cells remodels signaling components in recipient NoTf cells, including increased BMPR1b expression; (ii) identify BMPR1b as a receptor machinery transported within TNT, and as a specific regulator of TNT formation.

### After TNT contact with transformed donor cells, non-transformed acceptor cells become sensitized to a transforming microenvironment

First, by comparing our proteomic and bulk transcriptomic datasets [6], we identified three tumor-associated genes significantly upregulated during transformation: HOXB6 [28], CCND2 [29], and EREG [30] (Figure 7A). Their mRNA levels were increased in NoTf acceptor cells connected through TNTs with Tf cells, compared to NoTf cells in monoculture, and to NoTf acceptor cells for which TNT formation was hindered in Transwell (Figure 7B). Although these three genes may not be considered as direct drivers of a preneoplastic transformation, they could *bona fide* constitute relevant indicators of phenotypic changes, mediated through TNT cell-cell communication. As a negative control, we selected UFM1, a gene involved in endoplasmic reticulum stress, a cellular insult suggested to contribute to TNT development [31]. UFM1 transcript levels remained unchanged regardless of transformation status or material uptake (Supplementary Figure S9A-B), suggesting that although ER stress may be present, it is probably not a key driver of the cellular behavior described here.

**Figure 7.**
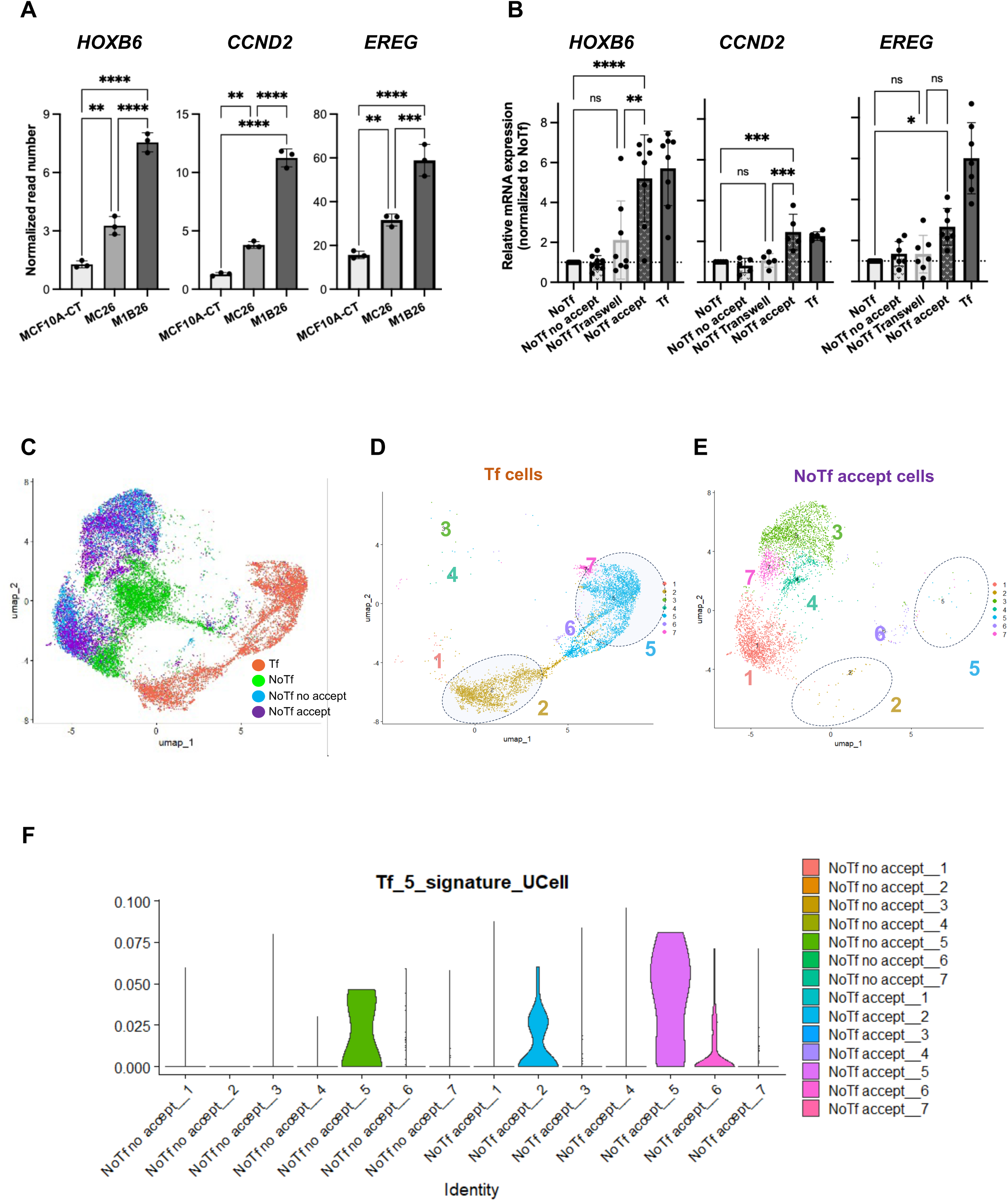
TNT-mediated transfer reprograms recipient cells toward a transformed-like transcriptional state. A, RNA-sequencing results of transcriptomic expression levels of HOXB6, CCND2, EREG and HRAS in indicated cell lines (n = 3 biological replicates). Data deposited on the Gene Expression Omnibus repository, RRID:SCR_005012, GSE186734 [6]. Error bars depict SD. Statistical significance was calculated using a one-way ANOVA with post-hoc Tukey’s Honest Significance Difference test (* p < 0.05; ** p < 0.01; *** p < 0.001; **** p < 0.0001). B, RT-qPCR measurements of HOXB6, CCND2, and EREG mRNA in indicated cell populations, with the expression in NoTf cells set at 1. Error bars depict SD. Statistical significance was calculated using a one-way ANOVA with post-hoc Tukey’s Honest Significance Difference test (biological replicates: n = 8 for HOXB6, n = 5 for CCND2, n = 7 for EREG). For clarity, only comparisons between NoTf, NoTf Transwell and NoTf accept cells are shown (* p < 0.05; ** p < 0.01; *** p < 0.001; **** p < 0.0001). C, U-maps comparing single cell transcriptomic profiles of Tf, NoTf, NoTf no accept and NoTf accept cells. D-E, U-maps at single cell resolution displayed as a Uniform Manifold Approximation and Projection (UMAP), based on identified clusters (7 different clusters) of D, Tf cells or E, NoTf accept cells. F, UCell signature scoring plots: Tf_5_signature_UCell corresponds to the 30 most highly expressed genes identified in cluster 5 of Tf cells. The plots show the distribution of this signature across clusters of NoTf cells, distinguishing NoTf accept from NoTf no accept cells.

Second, the transcriptomic heterogeneity associated with TNT-mediated material transfer was assessed by single-cell RNA sequencing of parental and sorted cells after direct co-cultures of NoTf with Tf cells (NoTf accept, NoTf no accept, Figure 4A). UMAP representation reveals that NoTf, NoTf accept, and NoTf no accept formed three distinct but closely related clusters (Supplementary Figure 9C), indicating that coculture conditions induced modest transcriptional changes in NoTf cells. To specifically assess the impact of material acquisition, we compared NoTf accept with NoTf no accept cells, along with Tf cells (Figure 7C). We thereby identified two sub-populations in NoTf accept cells which clustered within the transcriptional landscape of Tf cells, but distinct from NoTf no accept cells (Figure 7C). This indicates a shift of these recipient sub-populations toward a transformed-like transcriptional state. We then focused on the transformation-associated genes BMPR1B, HOXB6, CCND2 and EREG, which were specifically enriched at the single cell level in the NoTf accept cells subpopulations and Tf cells (Supplementary Figure 9D).

Single-cell UMAP analysis of Tf (Figure 7D) and NoTf accept cells (Figure 7E) unveiled seven distinct transcriptional clusters in each population. We identified cluster-specific gene signatures for the two most prominent Tf clusters (clusters 2 and 5) and used UCell scoring to assess their enrichment across the transcriptional clusters of NoTf accept and NoTf no accept cells. The Tf_5_signature_UCell, corresponding to the 30 most highly expressed genes in Tf cluster 5, was preferentially enriched in cluster 5 of NoTf accept cells (Figure 7F), with a lower but detectable enrichment also observed in cluster 2. Interestingly, a moderate signature enrichment was also detected in NoTf no accept cluster 5 cells. This suggests that, in addition to recipient cells undergoing post TNT-transfer transcriptional changes, a minor amount of NoTf cells may intrinsically display partial similarity to the Tf cluster 5 transcriptional program, consistent with the previously reported intrinsic heterogeneity of the MCF10A cell line [6]. In contrast, the transcriptional signature derived from the Tf cluster 2 (Supplementary Figure S9E) showed a weaker enrichment pattern across NoTf clusters. This indicates that cells displaying similarities to the Tf cluster 2 transcriptional profile are present at very low frequency within the NoTf population. Therefore, TNT-mediated transfer concurs with the emergence of recipient cell sub-populations with transcriptional features highly similar to Tf cells, while revealing the presence of rare pre-existing NoTf cells with partial similarity to Tf cluster 2 and 5 signatures.

Functionally, we first addressed whether TNT intercellular connections varied with cell cycle phases. This was investigated using the Fucci(CA) genetic tool, readily discriminating cells in various phases of the cycle by optical strategies [32]. Supplementary Figure S10A indicates that cells, transformed or not, could form TNTs at any stages of their cycle. Then, to unravel the potential impact of the differentiation and immaturity status of cells on TNT formation, we co-immunostained cells with CD10, a stem marker, Cytokeratin 14 (CTK14), a myoepithelial marker, and/or Cytokeratin 18 (CTK18), an epithelial marker [6]. Our findings suggest that any type of mammary cell population is capable of TNT formation, suggesting an absence of correlation between the differentiation and immaturity status, and the capacity of cells to form TNTs (Supplementary Figure S10B).

Then, the long-term impact of cell-cell communication was assessed on sorted cells, cultured for up to 16 weeks in the absence or presence of BMP2/IL6 (Figure 8A), thereby mimicking *in vivo* luminal breast tumor conditions, i.e. an inflammatory context, and the dysregulation of the BMP signaling pathway of mammary epithelial cells, due to high levels of BMP2 secreted by the tumor niche [4,33]. BMPR1b protein levels rose between weeks 6 and 10 before stabilizing, in NoTf accept cells exposed to cytokines after sorting (Figure 8B left), reaching the highest expression among sorted populations by week 14 (Figure 8B right; Supplementary Figure S11 A-C). In the tumor progression model we developed, recapitulating the clinical relevance of early steps of luminal breast tumor initiation and progression [3,5] (Figure 1A), enhanced BMPR1b expression correlated with the activation of canonical BMP signaling, as identified in vivo [4]. This was evidenced by enhanced phosphorylation of SMAD1/5/8, and of non–canonical MAPK, including p38 and ERK (Supplementary Figure S7A,B). After an 8-day TNT contact with Tf cells, followed by sorting and re-culture, NoTf accept cells displayed sustained activation of both p-SMAD1/5/8/SMAD1/5/8 and non canonical p-p38/p38 signaling pathways, whether BMP2/IL6 were present or not, with p38 activation peaking at 4 weeks post-transfer and canonical SMAD signaling reaching maximal activation at 6 weeks (Figure 8C-D; Supplementary Figure S11D-E). Finally, NoTf acceptor cells cultured in the presence of the pro–transforming cytokines BMP2 and IL6 [4,6] for up to 16 weeks following TNT transfer and cell sorting (Supplementary Figure S12A), exhibited anchorage-independent growth earlier than any other sorted cell populations, ultimately forming colonies in numbers comparable to those of Tf cells (Figure 8E; Supplementary Figure S12B). Potential cross-contamination between Tf and NoTf sorted cell samples was ruled out based on the specific fluorescence of derived soft agar clones (Supplementary Figure S12C). All these features are hallmarks of a preneoplastic transformation, observed exclusively in cells that had established TNT contacts with transformed cells prior to sorting, enabling the transfer of transforming information.

**Figure 8.**
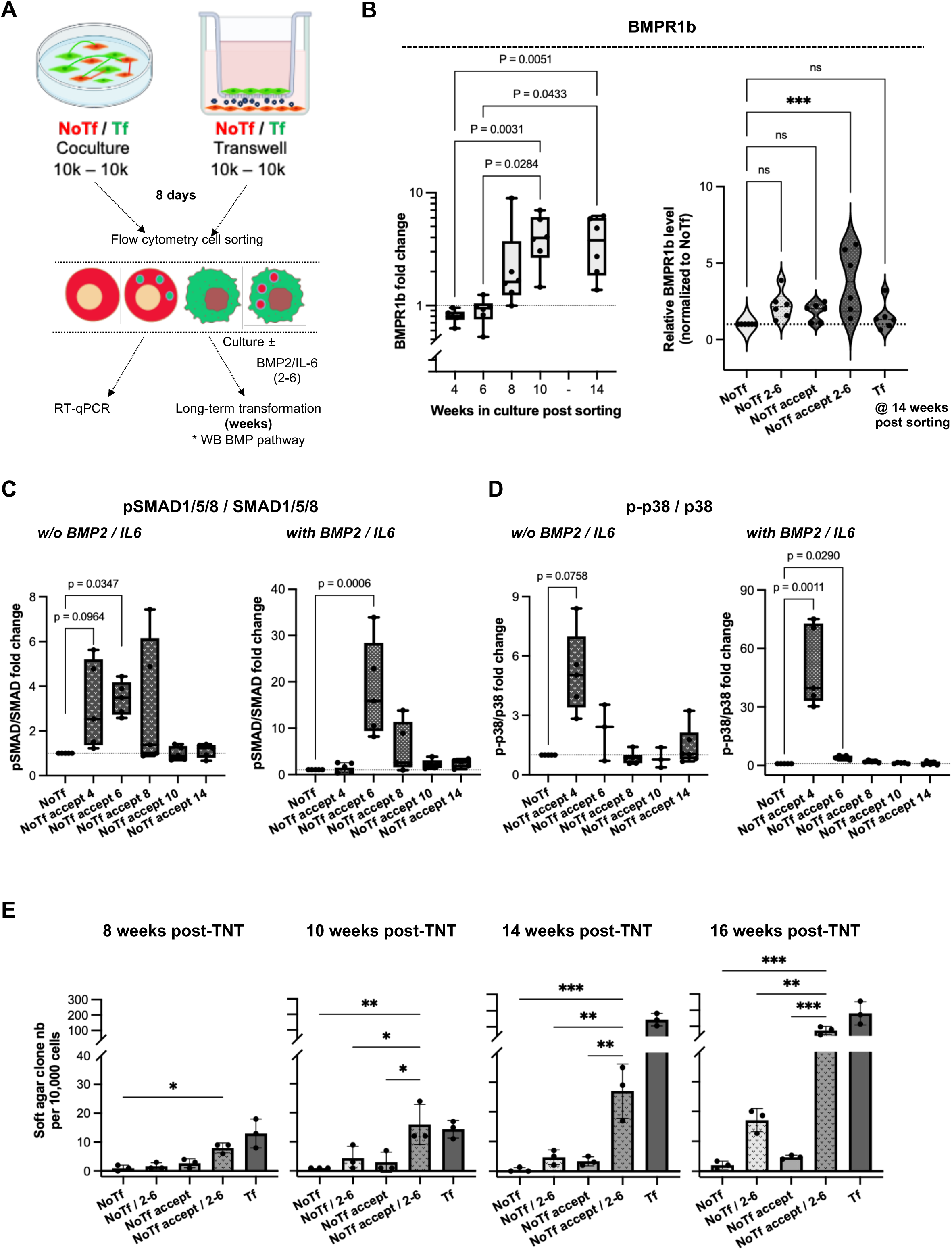
**Non-transformed cells acceptor of information from transformed cells *via* TNTs display preneoplastic features**. A, Schematic of experimental design: after sorting, cell populations were either lysed at day 8, and subjected to RT-qPCR for selected transcript analyses, or grown for several weeks with or without BMP2/IL6, and immunoblot analyses performed at end points. B, Left, quantification of BMPR1b immunoblots performed on NoTf accept cells grown in the presence of BMP2/IL6, for indicated weeks post sorting. Box and whiskers plot of 6 biological replicates. Statistical significance was calculated using a one-way ANOVA with non-parametric Kruskal-Wallis H test, and for clarity, only p-values ≤ 0.05 are indicated. Right, quantification of BMPR1b immunoblots performed on indicated cells, 14 weeks post sorting, with NoTf conditions set at 1 (n = 6 biological replicates). Statistical significance was calculated using a one-way ANOVA with post-hoc Tukey’s Honest Significance Difference test (*** p < 0.001). C-D, Quantification of pSMAD1/5/8/SMAD1/5/8 (C) and p-p38/p38 (D) immunoblots performed on NoTf accept cells grown in the absence (left panels) or presence (right panels) of BMP2/IL6, for 4, 6, 8, 10, or 14 weeks post sorting. Box and whiskers plot of 6 biological replicates. Statistical significance was calculated using a one-way ANOVA with non-parametric Kruskal-Wallis H test, and for clarity, only p-values ≤ 0.05 are indicated. E, Quantification of soft agar clones of indicated sorted cells, after 8, 10, 14 and 16 weeks in culture with or without BMP2/IL6 (n = 3 biological replicates). Error bars represent SD. Statistical significance was calculated using a one-way Welch and Brown-Forsythe ANOVA test (* p < 0.05; ** p < 0.01; *** p < 0.001). For clarity, only p-values ≤ 0.05 are indicated, and comparisons with Tf cells are omitted.

Hence, these findings unravel the key role of TNT-mediated transfer of transforming cues, including BMPR1b, in driving a preneoplastic switch in previously non-transformed cells receiving material from transformed cells. This TNT communication would prime non-transformed cells to respond to a tumor-promoting microenvironment (Figure 9).

**Figure 9.**
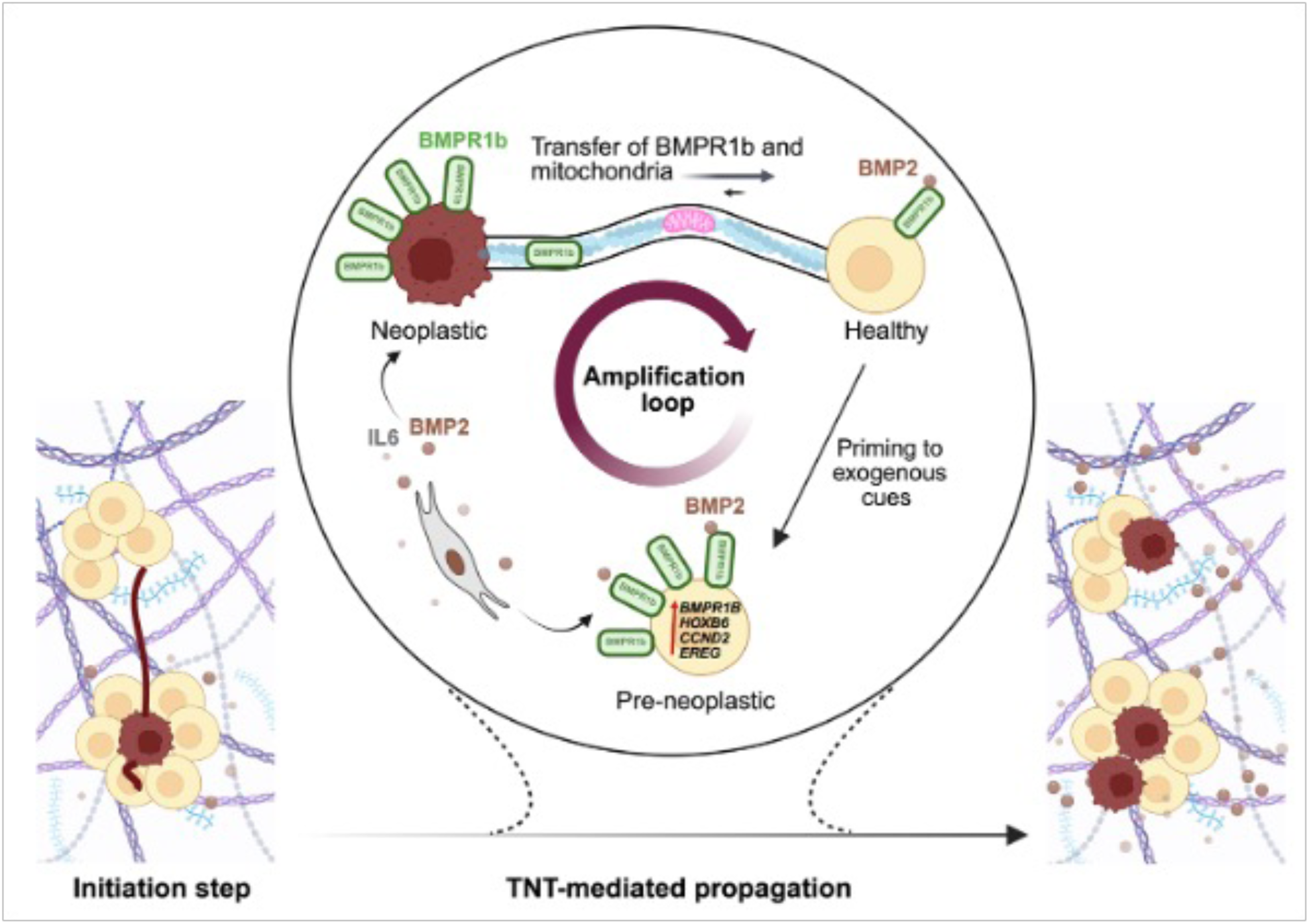
Model depicting the involvement of TNTs in the diffusion of preneoplastic cues involving the BMP signaling pathway. Fibrils and fibers illustrate extracellular matrix components.

## Discussion

Tunneling nanotubes (TNTs) have been extensively implicated in tumor progression, metastasis, and chemoresistance, through the intercellular transfer of mitochondria, notably from stromal cells to cancer (stem) cells [34,35], and more recently from neurons to cancer cells [13].

Building on this framework, our findings reveal that TNT–mediated transfer of transforming cues from transformed to non–transformed cells could activate BMP signaling in recipient cells, concomitant with increased BMPR1b expression, thereby sensitizing them to a tumor–promoting microenvironment and promoting the early acquisition of preneoplastic features.

In contrast, the delivery of material from transformed cells to non-transformed neighboring epithelial cells, remains unexplored at very early stages of transformation. Our study is the first to investigate the functional impact of TNT-mediated material exchange between cells of identical phenotype (i.e. epithelial cells), on the predisposition of non-transformed cells to adopt a preneoplastic status. Although donor-to-acceptor material transfer through secretion of soluble compounds and extracellular vesicles occurred, it remained low to negligible, indicating that the processes mainly relied on physical TNT-mediated communications. This horizontal transfer, occurring preferentially from transformed to non-transformed cells, leads within days to the overexpression of tumor-related genes and of BMPR1b itself, evocative of a possible feedback loop that would maintain transforming conditions (Figure 9). Non-transformed cells also undergo profound changes of their proteomic and transcriptomic landscapes, notably linked to transport processes, and within weeks acquire phenotypic traits of preneoplasia, such as anchorage-independent growth, BMPR1b over-abundance and related mounting of a non-canonical BMP-mediated signaling cascade. The earliest events, within one week, comprised the overexpression of tumor-related genes such as HOXB6, CCND2 and EREG by non-transformed acceptor cells, after TNT contacts with transformed (donor) cells. Interestingly, upregulation of these genes upon transformation has already been linked to other cancers: (i) HOXB6, encoding a homeobox domain-containing transcription factor involved in development, is upregulated in acute myeloid leukemia and osteosarcoma [28,36], with a possible role in cancer initiation [37]; (ii) CCND2, encoding cyclin D2, involved in DNA replication, cell division and cell cycle regulation, was dysregulated in gastric, thyroid and colorectal cancers [38], and its overexpression in breast cancer promotes MAPK-mediated cell proliferation and tumor growth [29,39]; (iii) upregulation of EREG, encoding the secreted protein ligand of the EGF receptor (EGFR) epiregulin, is linked to breast cancer progression and resistance to tamoxifen by EGFR activation [30,40]. Interestingly, experiments on TNT-forming capacities of mammary cell models, where transformation of either normal primary [23] or MCF10A cells [41] is « switched on » by the H-Ras oncoprotein, showed fewer and shorter TNTs connecting cells. HRAS is overexpressed in invasive breast tumors, but the prevalence of HRAS mutations in luminal types is negligible [1,42]. Although powerful, these mammary models do not faithfully recapitulate the earliest stages of non-invasive luminal breast tumors triggered, for example, by exposure to carcinogenic cues such as environmental pollutants, endocrine disruptors or alcohol [4,43–45].

Following material transfer in TNTs, including the BMPR1b receptor in their lumen, BMPR1B becomes overexpressed in acceptor cells within a week. The receptor molecule was more abundant in transformed cells, was detected within TNTs, contributed to their elongation, and was transferred to non-transformed cells through TNT-like structures. As BMP receptors are known to be located in lipid rafts, BMPR1b could induce TNT formation *via* its interaction with these microdomains [46], from which TNTs might arise [47,48]. These cholesterol-rich microdomains provide high curvature and elasticity necessary for membrane deformation linked to TNT formation. Additionally, BMP2 was reported to promote actin polymerization via PI3Kα activation [49]. We herein observed that LCP1 was upregulated upon transformation, concomitantly to its protein Plastin-2, in acceptor cells following TNT transfer. Plastin-2 is a key actin-bundling protein involved in the formation of cellular protrusions [50], overexpressed in epithelial-derived cancers including breast [51]. Its overexpression correlated with cancer cell migration, invasion, metastasis [52] and chemoresistance [53]. Plastin-2 activation depends on serine 5 phosphorylation *via* ERK/MAPK, PI3K, and PKA/PKC pathways, enabling its recruitment to actin filaments and promoting bundling and protrusion elongation, as shown in breast carcinoma [54]. This agrees with previous findings, showing that TNT induction by Eps8 involves its actin-bundling activity [55]. Altogether, this might explain the higher number of TNTs observed upon transformation.

At the transcriptional and functional level, the analysis of sorted acceptor cells which received material from TNT transfer, revealed their gradual transformation over weeks, faster and more extensively than non-acceptor cells, concomitant to a progressive but sustained activation of the BMP pathway. This induced an initial burst of the non-canonical BMP-mediated p38 downstream response, followed by the prolonged activation of the canonical SMAD cascade, only observed when TNT-like contacts existed. The p38 MAPK pathway is a well–established regulator of mammary epithelial cell transformation, acting as an early stress–responsive barrier through the induction of cell–cycle arrest, oncogene–induced senescence, and apoptosis [56], while being frequently bypassed or reprogrammed during tumor progression to support cellular plasticity [57]. Beyond its role in stress signaling, p38 also regulates cytoskeletal dynamics, lamellipodium formation and cell migration [49,58]. Consistent with this framework, our data are in favor of a TNT–mediated material transfer, which would trigger a rapid activation of the non-canonical BMP-mediated p38 pathway, which is then efficiently repressed, thereby enabling the emergence of preneoplastic traits in recipient cells.

Here, we unveil for the first time that TNTs transport BMPR1b and could spread transforming cues, in the context of luminal breast cancer initiation. Such processes sensitize acceptor cells to BMP2 signals of the microenvironment, rapidly triggering a cascade of molecular events that culminate in phenotypic changes, with anchorage-independent growth of previously non-transformed cells (Figure 9). Further studies using *in vivo* models will be valuable to assess the relevance of these findings in physiological and clinical settings. Nevertheless, combined with the recent demonstration that TNTs can transfer material *in vivo* [59], our work provides the foundation for a new concept: the propagation, through TNTs, of preneoplastic cues within a population of initially non-transformed mammary cells. This process could precede the development of luminal-type tumors, and identifies BMPR1b as a major early contributor of the BMP-dependent preneoplastic switch of acceptor cells.

## Materials and Methods

### Diversa

No samples size calculations were performed in our study. With the exception of human samples restricted by availability, the size of other experiments was obtained for a biological minimum for statistical applicability. All relevant samples available were provided for the study. No relevant data was excluded from this manuscript.

A minimum of three replicates were performed for each experiment helping the evaluation of reproducibility of data. All of the replicates are plotted and used in the evaluation of our hypothesis. No randomization nor blinding have been performed.

### Reagents

TNTi was a kind gift from Christel Vérollet, IBPS, Toulouse [20,21]. Cytochalasin D was purchased from Sigma Aldrich (#504776).

### Cell lines

All cell lines were monthly tested for mycoplasma contamination, by PCR through Eurofins Europe analysis. MCF10A cells were purchased from ATCC (ATCC Batch #7635052, RRID:CVCL_0598). The MC26 cell line was obtained upon MCF10A chronic exposure to BMP2 and IL6, while M1B26 came from MCF10A first sorted on BMPR1b overexpression, then chronically exposed to BMP2 and IL6, and derived from a soft agar colony [6]. MCF10A, MCF10A.DCIS, MC26 and M1B26 were cultured in phenol-red free Dulbecco’s Modified Eagle Medium Ham’s F12 (DMEM-F12, Gibco), supplemented with 2% horse serum (Gibco), 10,000 U/ml - penicillin and 10,000 μg/ml - streptomycin (Gibco) (P/S), 10 μg/ml insulin (NovoRapid CLB pharmacy), 0.5 μg/ml hydrocortisone (Sigma), 100 ng/ml cholera toxin (Sigma), 20 ng/ml epithelial growth factor (Sigma). MCF10A.AT1 and MCF10A.DCIS, obtained from ATCC, were cultured with the same medium supplemented with 5% horse serum. MC26 and M1B26 culture medium was supplemented with 5 ng/ml BMP2 (R&D) and 10 ng/ml IL6 (Peprotech). HMLE and HMLER cell lines were kindly provided by Robert Weinberg (MIT, Massachussetts, USA), under the MTA number CLB I-3991-01 (signed on 1/25/2024). Cells were grown in 1:1 MEGM Complete Medium [Lonza/Clonetics, CC-3150] plus DMEM/F12 (1:1) with supplements : 10 ng/ml hEGF, 0.5 μg/ml hydrocortisone, 10 μg/ml insulin, 10,000 U/ml - penicillin and 10,000 μg/ml - streptomycin (Gibco). Cells were passed twice a week at 70% of confluence. Cells washed twice with Phosphate Buffer Saline (PBS, Life Technologies) were treated with 0.48 mM Versene (ThermoFischer, #15040066) for 20 min, or with trypsin 0.05% for 10 min at 37°C. After collecting cells, neutralization was achieved with PBS/20% FBS. Plated cells were incubated at 37°C under 5% atmospheric CO_2_.

### Normal human primary mammary epithelial cells

Samples for normal human primary mammary epithelial cells and fibroblasts were obtained from healthy women undergoing mammary reduction, with no pathophysiological background and no age limit defined. Healthy breast samples obtention was approved by the CPP (Comité de protection de personnes-French ethics committee) number 20.01.31.72653 and 22.02667.000116. All patients gave informed consent. Human mammary epithelial and stromal cells were prepared from reduction mammoplasty, as described in [4].

### Fluorescent cell lines generation

Fluorescent cells were obtained by viral transduction of parental ones with lentivirus produced in 293T cells (RRID:CVCL_0063) by transfection of three plasmids encoding respectively for the viral envelope G of VSV (vesicular stomatitis virus), the structural and enzymatic proteins Gag and Pol of the lentivirus, and for a reporter gene mCherry or GFP (in collaboration with AniRA lentivectors production facility from the CELPHEDIA Infrastructure and SFR Biosciences, UAR3444/CNRS, US8/Inserm, ENS de Lyon/UCBL). Of note, these viruses are defective, meaning that no viral protein or particle are produced by cells after transduction.

### mCherry-tagged BMPR1b expression in GFP-labeled M1B26 cells

An expression plasmid construct for an mCherry-tagged version of human BMPR1b [pLV[Exp]-Puro-hPGK>hBMPR1B[NM_001256793.2]/mCherry (VB240328-1316ggp] was designed and purchased from Vector Builder (Chicago, USA), and delivered as *E. coli* stock. On the day of the transfection (day 0), 1 million GFP-labeled M1B26 cells were resuspended in 82 μl of the Lonza™ Nucleofector^®^ solution, and 18 μl of Lonza Supplement 1 containing 2 μg of purified plasmid DNA. After electroporation (Lonza™ Nucleofector^®^), cells were immediately transferred into complete warm M1B26 culture medium and grown for 24h. On day 1, puromycin (500 ng/ml final) was added to the culture medium. After a week, puromycin was omitted from the medium, and cells maintained in M1B26 medium supplemented with 5% horse serum.

### Coculture of MCF10A-CT and M1B26 cells

Cells were cocultured at a 50% ratio for up to 8 days in MCF10A complete medium. Fresh medium was added every 3-4 days. For the conditioned medium assay, the M1B26 green or MCF10A-CT green supernatant was collected, centrifuged for 5 min at 800xg, and filtered on a 20 µm-pore filter. Monocultures of fluorescent cells were grown for 8 days in this medium, and fresh one was added after 3-4 days. For cocultures in Transwell, a Transwell system with pores of 0.4 µm was used (Merck Millipore, ref PHP 01250). Four and ½ ml of MCF10A complete medium was added at the bottom, and 450 µl of MCF10A complete medium at the top of the insert. Cells were cocultured in the Transwell system for 8 days, and fresh complete medium was added after 3-4 days.

### Acini formation

The protocol for 3D-acini formation was derived from Vidi *et al* [60], using the 3D-embedded culture process.

### Gene silencing with siRNA

Silencing of BMPR1b and GPER was performed using small interfering RNAs, purchased at Horizon Discovery (ON-TARGETplus siRNA, SMARTpool siRNA). Cells were transfected with 100 nM siRNAs, through electroporation by a Neon™ Electroporator Device, and grown for 48h (siBMPR1b) or 72h (siGPER). Fresh medium devoid of siRNAs was then added. ON-TARGETplus Non-targeting Pool of siRNAs were used as controls. For some experiments, M1B26 cells were treated with 50 nM siRNA targeting BMPR1b, or with non-targeting siRNA for 24h, supplemented with an additional 50 nM for 12h. Cells were then used for appropriate controls (viability, silencing yield, western blot)) and for further experiments.

### Migration assay

Cells plated in 96-well plates at high confluency (50,000 cells/100μl) were grown overnight in complete medium. The following day, after PBS washing, cells were treated with Mitomycin C from Streptomyces caespitosus (Merck Sigma, #M0503), at a final concentration of 5μg/ml for 2h at 37°C, to prevent cellular division. After PBS washing, a scratch was performed, using the scratch wound plate. After PBS washing to eliminate detached cells, cells were left at 37°C for 1h, and imaged during 24h by Incucyte^®^ Birdo microscopy.

### Invasion assay

Cells were plated at high confluency (50,000 cells/100μL) in a 96-well plate coated with phenol red free-Matrigel (100μg/ml Matrigel, 50μl *per* well), and left overnight in complete medium. The following day, after PBS washing, cells were treated with Mitomycin C from Streptomyces caespitosus (Merck Sigma, #M0503) at a final concentration of 5μg/ml for 2h at 37°C, to prevent cellular division. After PBS washing, a scratch was performed using the scratch wound plate. After PBS washing to eliminate detached cells, 50μl of Matrigel (1/25 dilution) were added, and cells left at 37°C for 1h. This was followed by imaging 24h by Incucyte^®^ Birdo microscopy.

### Western-Blotting

Cells were harvested, pelleted, and lysed in CHAPS buffer (40 mM Hepes pH 7.4, 120 mM NaCl, 2 mM EDTA, 0.3% CHAPS) and protease/phosphatase inhibitors (Roche Complete inhibitor cocktail #4693132001). Proteins were then assayed by spectrophotometry, along with a BSA standard curve, using the Pierce™ Coomassie Protein Assay Kit (Thermo Fisher Scientific). Twenty to 30 μg of proteins were used and mixed with 1X Laëmmli and H_2_O, and samples boiled for 5 min at 95°C. The mixture was loaded onto 10% acrylamide gels, and run in 25 mM Tris, 192 mM glycine, 0.1% SDS. After transfer onto PVDF membranes previously activated in ethanol and transfer buffer (Biorad™ Transfer buffer, 20 % ethanol) using the Transblot Turbo^TM^ from BioRad. Membranes were then incubated for 1h with TBST [Tris-buffered saline 1X (Euromedex) containing 0.1% Tween-20] with 5% fat-free BSA for membrane saturation at room temperature. All antibodies used are commercially available and tested by manufacturers. All antibodies stainings were done in parallel with negative controls to guarantee specificity. Primary antibodies were incubated at 4°C overnight. Primary antibodies used were directed against BMPR1b (1:300 Abcam, ab78417 RRID:AB_1565836), GAPDH (1:50,000, Proteintech #60004-1-Ig RRID:AB_2107436), p-SMAD1/5/8 (1:500, Cell signalling, #13820), Smad1/5/8 (1:500, Santa Cruz, sc-6031-R), p-ERK (1:2,000, Cell Signalling Technology, #4370S), ERK (1:1,500, Cell Signalling Technology #4695S), p-p38 (1:2,000, BD Bioscience #612288), p38 (1:2,000, BD Biosciences, #612168), GPER (1:500 Novus Biologicals, #NLS4290 or 1:500, Invitrogen, #PA5-28647), or HoxB6 (1:1,500, Santa Cruz, sc-166950), in 5% BSA. After TBST buffer washes, membranes were incubated at room temperature for 1h with horseradish peroxydase (HRP)-conjugated secondary antibodies goat anti-rabbit, goat anti-mouse or rabbit anti-goat secondary antibodies (Jackson, #111-035-003, #115-035-003, #305-035-003) at 1:10,000 in TBST/3% BSA. After extensive washings with TBST buffer, revelation was achieved through Clarity™ Western ECL Substrate (Biorad) and Chemidoc™ Imaging System (Biorad). Protein bands were quantified using the ImageLab™ software.

### Immunofluorescence

Cells were seeded on coverslips and grown at 60% of confluence. For BMPR1b and actin staining, the following protocol was used : after washing with PBS, fixation was performed for 20 min in 4% final paraformaldehyde (PFA) at room temperature (RT). After PBS washing, permeabilization and saturation were performed for 45 min with PBS/3% BSA/0.3% Triton X-100 at RT. After PBS washing, primary antibody against BMPR1b (1:300 #sc-293428 Santa Cruz) and its corresponding isotype were incubated overnight at 4°C. Coverslips were washed three times with PBS/3% BSA/0.1% Tween-20 for 10 min. Secondary antibody goat anti-mouse Dylight 488 diluted 1:1,000 was incubated for 45 minutes at RT, with 1:50,000 of Hoechst 33342 (Invitrogen #H3570) and Phalloidin AlexaFluor-Plus 647 (Invitrogen, #A30107). Then, cells were washed twice with PBS/3% BSA/0.1% Tween-20 for 5 minutes and twice with PBS for 5 minutes. Coverslips were then mounted using the Dako mounting medium (Agilent, ref S3023). For mitochondria staining, MitoTracker^®^ deep red or green were used (Thermofischer, #M22426 or #M7514, respectively). For some experiments, GFP-labeled M1B26 cells expressing an mCherry-tagged BMPR1b construct, fixed and permeabilized as described above, were first incubated with a fluorescence signal enhancer Image-iT™ FX (ThermoFisher™, #I36933), according to manufacturer’s instructions. Then staining with a primary antibody against mCherry (mCherry E5D8F Rabbit monoclonal, Cell Signaling Technology™, #43590, 1:200 dilution) was performed, followed after washings by incubation with a secondary goat anti-rabbit AlexaFluor^®^647 (1:1000 dilution). Microscopic observations were performed with the objective 40x oil of a confocal Leica Stellaris 5 microscope. This was performed at the PIC platform of the CRCL, Lyon, France. Images were taken in 16-bit, and formatted *via* the Fiji software.

Other microscopic observations were performed using the x40 oil immersion objective of a confocal Zeiss LSM980 microscope, with Airyscan mode. This was done at the LYMIC-PLATIM, Imaging and Microscopy Core Facility of SFR Biosciences (UMS3444), Lyon, France. Images were taken in 16-bit, and formatted via the Fiji software (RRID:SCR_002285). For immunofluorescence staining of actin (Phalloidin) and tubulin, the following protocol optimized by the Dr. C. Zurzolo’s team was used to identify and characterize TNTs. Subconfluent cells were fixed for 15min at RT, with a 4% PFA/0.05% glutaraldehyde mixture in PBS. The fixation buffer was removed, and and cells incubated with 100 mM NH_4_Cl for 10min at RT to quench the PFA-related autofluorescence. After PBS washing, cells were saturated with 2% BSA for 20 min at RT. The primary antibody against tubulin was diluted at 1:500 (Abcam #ab6046, RRID:AB_2210370) in 2% BSA, and incubated for 1h at RT, or OVN at 4°C. After PBS washing, cells were incubated with a secondary goat anti-rabbit antibody-488nm, diluted at 1:1,000 in 2% BSA, and incubated for 40 min in the dark at RT. After PBS washing, cells were finally incubated with phalloidin AlexaFluor Plus-647 (Invitrogen, #A30107) diluted at 1:1,000, and with Hoechst 33342 (Invitrogen, #H3570), diluted at 1:50,000 in 1% BSA, for 30 min in the dark. After PBS washing, slides were mounted using the Dako mounting medium (Agilent, S3023). Microscopic observations were performed with the objective 40x oil of a confocal Zeiss LSM980 microscope, with Airyscan mode. This was performed at the LYMIC-PLATIM, Imaging and Microscopy Core Facility of SFR Biosciences (UMS3444), Lyon, France, and at the imaging quantitative center of Lyon Est (CIQLE). Images were taken in 16-bit, and formatted *via* the Fiji software. TNTs were finally manually counted with the Fiji software.

### Live imaging

Live imaging was performed after 48h of coculture in µslide 8-well plate with thin glass bottom (Ibidi, #80827), or in PhenoPlate™ 96-well microplates with optically clear flat bottom (Revvity, # 6055302), both adequate for imaging. For some experiments, after PBS washing, cells were stained with Cell Mask Deep Red (at 1:500, Invitrogen, #C10046), and Hoechst (at 1:500, Invitrogen, #H3570), diluted in complete medium, for 10 min at 37°C. After PBS washing, complete fresh medium was added. Observations were performed using the Opera Phenix HCS microscope with the 20x or 40x water immersion objectives.

For some experiments on BMPR1b-silenced GFP-labeled M1B26 cells, cocultures of these cells with unlabeled MCF10A-CT cells were stained for mitochondria tracking with Mitotracker^®^-deep red (647 nm), and imaged using the Opera Phenix HCS microscope.

### Extracellular vesicles isolation

For these experiments, cells were grown in serum-deprived medium, in the absence or presence of GW4869 at 20 μM [22]. Collected supernatants were sequentially centrifuged 10 min at 300xg, 30 min at 2,000xg, 30 min at 10,000xg, and then 90 min at 100,000xg. The pellet was then washed with PBS, and ultracentrifuged for 90 min at 100,000xg. The resulting pellet containing extracellular vesicles was resuspended in PBS containing the cOmplete^®^ protease inhibitor cocktail (Roche™, # 11697498001).

### RT-qPCR

For RT-qPCR analyses, pelleted cells were resuspended with 300 µl Trizol reagent (Life Technologies^TM^). Total RNA purification was achieved using direct-zol RNA MicroPrep w/ Zymo-Spin IC Columns extraction kit (Ozyme). For sorted fluorescent cells after coculture, RNA extraction was performed using the AllPrep DNA/RNA Micro Kit (Qiagen, #80284). C-DNA was then obtained using PrimeScript™ RT Reagent Kit (Takara kit), and qPCR then performed using the SyberGreen Premix Ex Taq™ (Takara kit) as the fluorescent DNA intercalant, and amplicons read by a Light Cycle 480 (Roche, LC480). Results were normalized to housekeeping genes : TATA-binding protein (TBP) and hypoxanthineguanine phosphoribosyl transferase (HPRT) (see Supplementary Table S4 for primers used). An additional normalization to control cells (MCF10A or MCF10A-CT) was also performed.

### Proteomic analyses

*Sample preparation for proteome analysis*. Proteome analyses were performed at the Prot’ICO proteomics facility: FACS-sorted samples (FACS-Melody) were quickly thawed and proteins equivalent to 66,000 cells were concomitantly extracted and denatured while thiol groups were chemically reduced in 46 µl 5% SDS, containing 5 mM TCEP, 10 mM chloroacetamide, heated in a thermostatic shaker set at 95°C under 500xg for 30 min. Sonication was performed twice 30s using an ultrasonic sonicator bath (ThermoFisher Scientific). S-Trap Micro Spin Column was used according to the manufacturer’s protocol (Protifi, Farmingdale, NY, USA), to trap precipitated proteins under acidic and hydrophobic conditions to remove detergents, and 1µg of porcine trypsin (Sciex) was used at 37°C overnight to cleave peptide bonds at carbonyl-side of lysine and arginine to generate bottom-up LC MS-compatible peptides. Resulting peptides were then cleared from potential insoluble material by centrifugation. Peptides were then speed-vacuum dried and resuspended in 0.1% formic acid (FA) before injection in LC-MS/MS with DIA-PASEF mode using a TIMS-TOF Pro 2 mass spectrometer (Bruker, Bremen, Germany).

*LC-MS/MS setup*. Each sample was loaded and separated on a C18 reverse phase column (Aurora Ultimate series 1.7µm particles size, 75µm inner diameter and 25cm length, IonOptics, Fitzroy, Australia) using a NanoElute 2 LC system with Hystar 6.3.1.8 software (Bruker, Bremen, Germany). Eluates were electrosprayed for a 60 min-long duration of the hydrophobicity gradient ranging from 99% of solvent A containing 0.1% FA in milliQ-grade H_2_O, to 27% of solvent B containing 80% acetonitrile containing 0.1% FA in mQ-H_2_O. The mass spectrometer acquired data throughout the elution process and operated in data-independent analysis (DIA) with PASEF-enabled method, using the TIMS-Control software v.5.1.8 (Bruker). Samples were injected in batch replicates in order to circumvent possible interferences of technical biases. The m/z acquisition range was 250–1201 with an ion mobility range of 0.7–1.25 1/K0 [V s/cm^2^], which corresponded to an estimated cycle time of one second. DIA-PASEF windows, and collision energy were also left to default with a base of 0.85 1/K0 [V s/cm^2^] set at 20 eV and 1.30 1/K0 [V s/cm^2^] set at 59 eV. TOF mass calibration and TIMS ion mobility 1/K0 were performed linearly using three reference ions at 622, 922, and 1,222 m/z (Agilent Technology, Santa Clara, CA, USA)

*LC-MS/MS raw data analysis*. The raw data were extracted, normalized and analyzed using Spectronaut® 19.0.24 (Biognosys A.G. Schlieren, Switzerland) in DirectDIA+ mode, which modelized elution behavior, mobility and MS/MS events based on the “Uniprot/SwissprotKB model organism_9606_2024_04_03” FASTA sequence database (RRID:SCR_011819) of 45,514 human proteins and 381 entries of common contaminant proteins. Protein identification false discovery rate (FDR) was restricted to 1% maximum, and inter-injection data normalization. The enzyme’s cleavage specificity was that of trypsin. The precursor and fragment mass tolerances were set to default. Acetyl (protein N-term) and oxidation of methionines were set as variable modifications, while thiol groups from cysteines were considered completely alkylated by carbamidomethylation. A minimum of two ratios of peptides was required for relative quantification between groups. Protein quantification analysis was performed using Label-Free Quantification (LFQ) intensities. Spectronaut also linked Gene Ontology terms to identified protein using the EBI GO database, and its statistical tools were used to visualize data, assess reproducibility and quality to allow quantitative dataset comparisons. The mass spectrometry proteomics data have been deposited to the ProteomeXchange Consortium via the PRIDE [61,62] (RRID:SCR_012052) partner repository with the dataset identifier PXD060023. Volcano plots were generated with the package EnhancedVolcano (https://bioconductor.org/packages/release/bioc/html/EnhancedVolcano.html), with an adjusted p-value (with Benjamini Hochberg procedure) threshold set at 0.05 and a Log_2_FC threshold set at 0.8. Pathway analysis were performed using STRING (https://string-db.org/; RRID:SCR_005223), with the Biological Process category from the Gene Ontology collection. For the search of BMP and p38 pathways targets, a combination of several pathways from different collections have been compiled (GOBP_POSITIVE_REGULATION_OF_BMP_SIGNALING_PATHWAY ; GOBP_REGULATION_OF_BMP_SIGNALING_PATHWAY ; GOBP_RESPONSE_TO_BMP ; PID_BMP_PATHWAY ; BIOCARTA_P38MAPK_PATHWAY ; GOBP_P38MAPK_CASCADE ; GOBP_REGULATION_OF_P38MAPK_CASCADE ; REACTOME_P38MAPK_EVENTS ; WP_P38_MAPK_SIGNALING).

### Single cell RNA sequencing analyses

FACS-sorted cell samples (FACS-Melody) were fixed with PFA 4%. Each sample was incubated with a specific Human WTA probes targeting the entire transcriptome. Samples were counted with a Luna-FL Dual fluorescence cell counter (Logos Biosystems), pooled together, washed, loaded on Next GEM Chip Q to obtain an expected cell recovery population of 3,000 cells *per* samples and run on the Chromium iX system (10X Genomics) according to manufacturer’s instructions. Gene Expression library was generated with the Chromium Single Cell Fixed RNA kit and sequenced on the NovaSeq 6000 platform (Illumina) to obtain around 50,000 reads *per* cell. After demultiplexing, data were processed using cellranger pipeline to generate cell count matrix for each sample.

*Data processing*. Raw count matrices were generated by Cell Ranger (10x Genomics) from 5 samples (M1B26_green, MCF10A_CTred, MCF10A_Ctred_Co_M1B26green, MCF10A_Ctred_Trans_M1B26green, MCF10A_CTred_green_from_M1B26green_Co). Ambient RNA contamination was estimated using decontX (celda package) with empty droplets from the raw feature-barcode matrix used as background. Cells with a contamination score above 0.75 were excluded. Raw counts were retained for downstream analyses. Doublets were detected and removed using scDblFinder.

*Quality control*. Cells were retained if they met the following criteria: between 200 and 8,000 detected genes, between 500 and 60,000 UMI counts, fewer than 20% mitochondrial reads, and a transcriptomic complexity (log10 genes / log10 UMI) above 0.8. Per-sample threshold overrides were applied for: M1B26_green (max_features = 8,000, max_mt_percent = 40); MCF10A_CTred (max_features = 5,000, max_mt_percent = 50); MCF10A_Ctred_Co_M1B26green (max_features = 8,000, max_mt_percent = 30); MCF10A_Ctred_Trans_M1B26green (max_features = 8,000, max_mt_percent = 40); MCF10A_CTred_green_from_M1B26green_Co (max_features = 8,000, max_mt_percent = 50).

*Normalisation and dimensionality reduction*. Gene expression counts were log-normalised (NormalizeData, scale factor 10,000). The top 3,000 highly variable genes were selected for dimensionality reduction. Principal component analysis (PCA) was performed and the first 20 principal components were retained for downstream analyses, as determined by elbow plot inspection. UMAP was computed for two-dimensional visualisation.

*Batch correction*. No batch correction was applied.

*Clustering and marker identification*. Cells were clustered using the Leiden algorithm at a resolution of 0.2 (resolutions 0.2, 0.4, 0.6, 0.8, 1, 1.2 were tested; final resolution selected based on clustree stability analysis). Cluster marker genes were identified using a Wilcoxon rank-sum test (Seurat FindAllMarkers; min.pct = 0.1, log2FC threshold = 0.25, positive markers only).

*Cell type annotation*. Automated cell type annotation was performed using SingleR with the following reference datasets: HumanPrimaryCellAtlasData and BlueprintEncodeData (label.main). Pruned labels were used to flag low-confidence predictions. Pan-Human Azimuth annotation was performed using the cloud API (Satijalab), providing hierarchical cell type labels across three levels of granularity. Final cell type labels were assigned manually based on automated annotation results, cluster marker genes, and expression of canonical cell type markers.

### Flow cytometry and cell sorting

After cell detachment as previously described, cells were resuspended in PBS/1% FBS. Sorting of cocultured cells was performed using the BD FACSMelody™ Cell Sorter, at the CRCL Flow cytometry platform, SFR Lyon-EST (CYLE). Results were analyzed with the FlowJo™ software.

### Soft agar colony formation assay

Each well of a 6-well plate was coated with agarose 1.5% mixed with 2X complete medium. After polymerisation at RT, a top layer was added, containing agarose 0.9%, mixed with 2X complete medium containing 13,000 cells. PBS was added between wells, so that the agar would not dry out. Clone counting was performed manually after 3-4 weeks.

### Electron microscopies

*Transmission EM.* Cell samples in Petri dishes were washed 3 times 15min in sodium cacodylate buffer 0.2M pH 7.3, and fixed for 1h in glutaraldehyde 2% in the same buffer at 0.1M. After washing in buffer, samples were prepared for TEM observations after Epoxy resin embedding and microtome ultrathin sectioning at the Center for Microstructure Technology (CTµ, Univ Lyon 1). Observations were performed with a Jeol 1400 Flash microscope operated at 200 kV, and equipped with a Gatan Rio 16 digital camera.

*Scanning EM.* SEM was performed in collaboration with the Center for Microstructure Technology (CTμ, Univ Lyon 1). Cells were fixed with glutaraldehyde 2.5% at RT diluted in phosphate buffer 0.1M, pH 7.2 for 3h. After 3 washings of 20 min with phosphate buffer 0.2M pH 7.2, cells were incubated with OsO4 1% final in phosphate buffer 0.1M pH 7.2 for 45 min. After H_2_O washings, cells were incubated overnight with 20% acetone in H_2_O. In a vacuum bell jar, a bowl with calcium chloride granules was placed in the lower part, and a glass petri dish with calcium sulfate granules in the upper part. Samples without lid were then introduced with enough quantity of 20% acetone, and vacuum applied for 24h. Acetone 100% was then added for 1h, and a critical point drying was performed for 2h. Cell imaging was performed using a scanning electron microscope Zeiss Merlin Compact VP.

*Correlative light scanning electron microscopy (CLEM).* CLEM was performed in collaboration with the Center for Microstructure Technology (CTµ, Univ Lyon 1). Cells were plated on standard square coverslips ITO (22x22mm), and fixed with PFA 4% 20 min. After three washes with phosphate buffer 0.2M pH 7.2 for 20 min, saturation was achieved using phosphate buffer with 3% BSA and 0.3% Triton X-100 for 45min. Incubation with primary antibodies and washings were performed as mentioned in the section “Immunofluorescence”. AlexaFluor-488 Fluoronanogold (FluoroNanogold™, Nanoprobes, #7201) was diluted at 1:100 in phosphate buffer with 1% BSA and 0.2% Tween-20, and incubated for 1h at RT. After washings, coverslips were mounted on a “Shuttle and Find” Zeiss device, and the observation was done with a Zeiss confocal LSM800 microscope. After confocal observation, coverslips were washed three times with H_2_O. Then silver enhancement was performed, using the HQ silver enhancement kit (Nanoprobes, #2012) for 5 minutes. After H_2_O washings, cells were incubated with 1% final OsO4 in PBS 0.1M pH 7.2, for 45min. After H_2_O washings, cells were incubated overnight with 20% acetone in H_2_O. The following day, 100% acetone was added for 1h, followed by critical point drying for 2h. SEM imaging was then performed at the Center for Microstructure Technology (CTµ, Univ Lyon 1), using a scanning electron microscope Zeiss Merlin Compact VP.

#### Statistical methods

Normal distribution was evaluated using the Shapiro-Wilk test. Mean comparisons of two samples were performed with an unpaired two-sided Welch’s t-test. One-way Anova with post-hoc Tukey’s Honest Significance Difference test, or with Dunnett’s correction test, or with non-parametric Kruskal-Wallis H test, or a one-way Welch and Brown-Forsythe ANOVA were used, when multiple comparisons were performed. Statistical analyses were performed using GraphPad Prism software (La Jolla, CA; RRID:SCR_002798). Significant p values are indicated by asterisks in the figures (* p<0.05 ; ** p<0.01 ; *** p<0.001, **** p<0.0001). For proteomics data, statistical analyses were carried out using R version 4.2.0 (The R Foundation for Statistical Computing) on RStudio 2022 (RStudio, Inc., Boston, MA). Data pre-processing was carried out using the R packages NOISeq and Caret, to filter features which were not statistically relevant from the dataset, normalise the data, impute zero values, and assess and remove sources of unwanted technical variation. Differential expression analysis was carried out using the R package Limma. The results were filtered for proteins with Benjamini-Hochberg FDR-adjusted p-values < 0.05 and Log_2_(FC) > 0.8. STRING (Search Tool for the Retrieval of Interacting Genes/Proteins; https://string-db.org) v12.0 was used for functional protein association network analysis to predict protein-protein interactions.

## Data availability statement

The mass spectrometry proteomics data have been deposited to the ProteomeXchange Consortium via the PRIDE (https://www.ebi.ac.uk/pride/) [61,62] partner repository, with the dataset identifier PXD060023. Transcriptomic data have been deposited on the Gene Expression Omnibus repository, RRID:SCR_005012, GSE18673411. Single cell RNA sequencing data have been deposited on the Gene Expression Omnibus repository, RRID:SCR_005012, GSE343015.

## Acknowledgements

Single cell-RNA sequencing was performed by the genomic platform of the CRCL, supported by the SiRIC-LyriCAN program (grant INCa-DGOS-INSERM-ITMO cancer_18003). We thank C. Degletagne (Genomic platform, CRCL) for expert advise and technical contribution to single cell-RNA sequencing. Microscopy and image analyses were performed at the microscopy facilities of the CRCL (platform PIC), of the SFR Santé Lyon Est (platform CIQLE), and of the ENS Lyon (platform LYMIC-PLATIM, UAR 3444 / US8 Univ Lyon 1). We thank M. Gautier, C. Vanbelle, D. Ressnikoff, E. Chatre and J. Brocard for their expert technical contribution to microscopy experiments. Cell sorting was performed at the flow cytometry facility of the CRCL, and we thank P. Battiston-Montagne for her expert contribution. We thank the AniRA lentivectors production facility of the CELPHEDIA Infrastructure and SFR Biosciences (UAR3444/CNRS, US8/Inserm, ENS de Lyon, UCBL), especially G. Froment, D. Nègre and C. Costa.

We thank P. Mehlen, P. Juin, R. Tomasini, C. Vérollet and B. Bartosch for highly inspiring discussions, and B. Manship for english proofreading. Some illustrations were created using BioRender (https://BioRender.com).

This work was supported by Agence Nationale de la Recherche (VMS, ANR-10-LABX-0061, ANR-CESA-018-04 and Convergence PLAsCAN ANR-17-CONV-0002); Ligue Nationale contre le Cancer (CCM, doctoral grant), Dept Ain, Rhône (VMS), Saône et Loire (EIP) ; Fondation ARC (CCM, 4^th^ year doctoral grant; VMS, SFI20111203500, PJA20171206331) ; the ERiCAN program of Fondation MSD-Avenir (VMS/EIP, DS-2018-0015) and Associations [Déchaîne Ton Cœur (VMS); Ruban Rose Prix Avenir 2021 (VMS) ; Comité féminin pour le dépistage du cancer du sein 74 (VMS) ; Symatese Fondation (VMS), and Fondation pour la Recherche Médicale (équipe FRM EQU202203014695 VMS).

## Author Contributions

E-I.P. and V.M.S. designed and supervised the study, conceived methodological developments and acquired funding. C.C.M. and E-I.P. generated the proteomic and scRNA seq data. C.C.M. and E-I.P. designed and performed all microscopy experiments on living cells. C.C.M., T.T.N., A.B. and E-I.P. performed and interpreted the experiments. C.C.M., S.L. and B.G. collaborated in study design, contributed to methodological developments and data interpretations. K.G. performed the bioinformatic analyses of the proteomic data. L.T. and K.G. analysed the single cell RNA-seq data. C.D. and E-I.P. designed and performed SEM and CLEM microscopy experiments. F.G., A.B. and C.H. performed proteomic analyses, and F.G. interpreted proteomic data. E.D. provided primary breast samples, and assisted with their processing. C.C.M., V.M.S and E-I.P. wrote the manuscript. E-I.P. wrote the revised version of the manuscript, critically proofread by S.L. All authors reviewed the manuscript and approved the final version for publication.

