## Supplementary Figures and Tables for "Tunneling nanotubes propagate preneoplastic cues in a BMP-dependent manner"

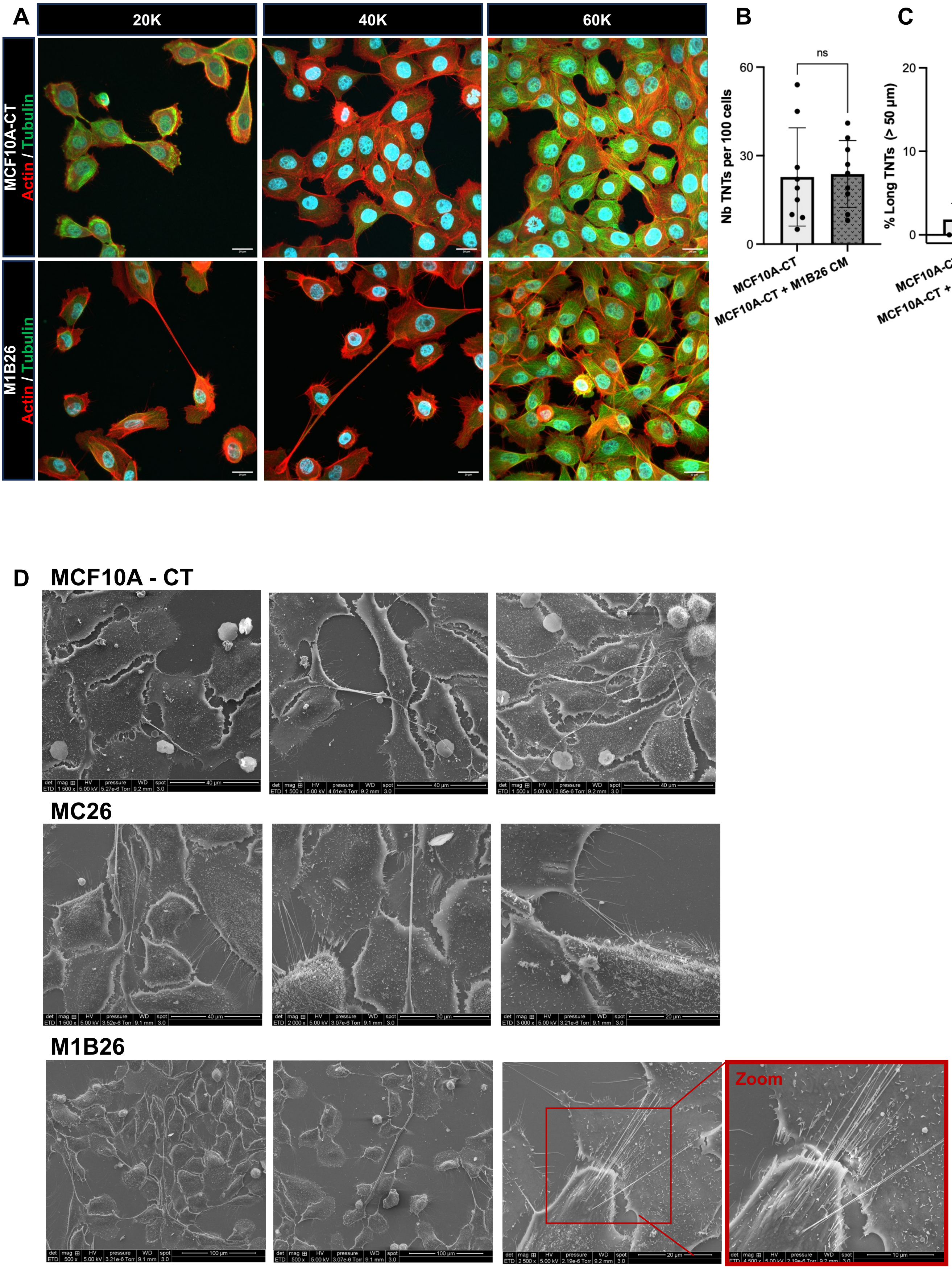


**Supplementary Figure S1**

**Supplementary Figure S1**. A, Representative immunofluorescence images of actin (red) and tubulin (green) in MCF10A-CT and M1B26 cells, plated at sub-confluent cell densities (20, 40 and 60K, respectively 20,000; 40,000 or 60,000 cells/$cm^2$), and observed by Airyscan confocal microscopy. Nuclei, Hoechst labeling (blue). Scale bars, 20 µm. B,C, Quantification of TNT number *per* 100 cells (B, n = 9 biological replicates), and % of long TNTs (C; > 50 µm; n = 4 biological replicates) for indicated cell conditions (MCF10A-CT+M1B26 CM : MCF10A-CT cells incubated for 48h with conditioned medium of M1B26 cells). Ten Z-stacks of 10 sections each were taken *per* condition, and 100 cells minimum *per* experiment were counted. TNTs were counted and measured in each Z-stack. Error bars depict SD. Statistical significance was calculated using an unpaired two-sided Welch's t test (B,C), and ns is not significant. D, Scanning electron microscopy images of TNTs of MCF10A-CT, MC26 and M1B26 cells. Scale bars MCF10A-CT: 40 µm; MC26: from left to right: 40, 30, 20 µm; M1B26: from left to right: 100, 100, 20; in zoom: 10 µm.

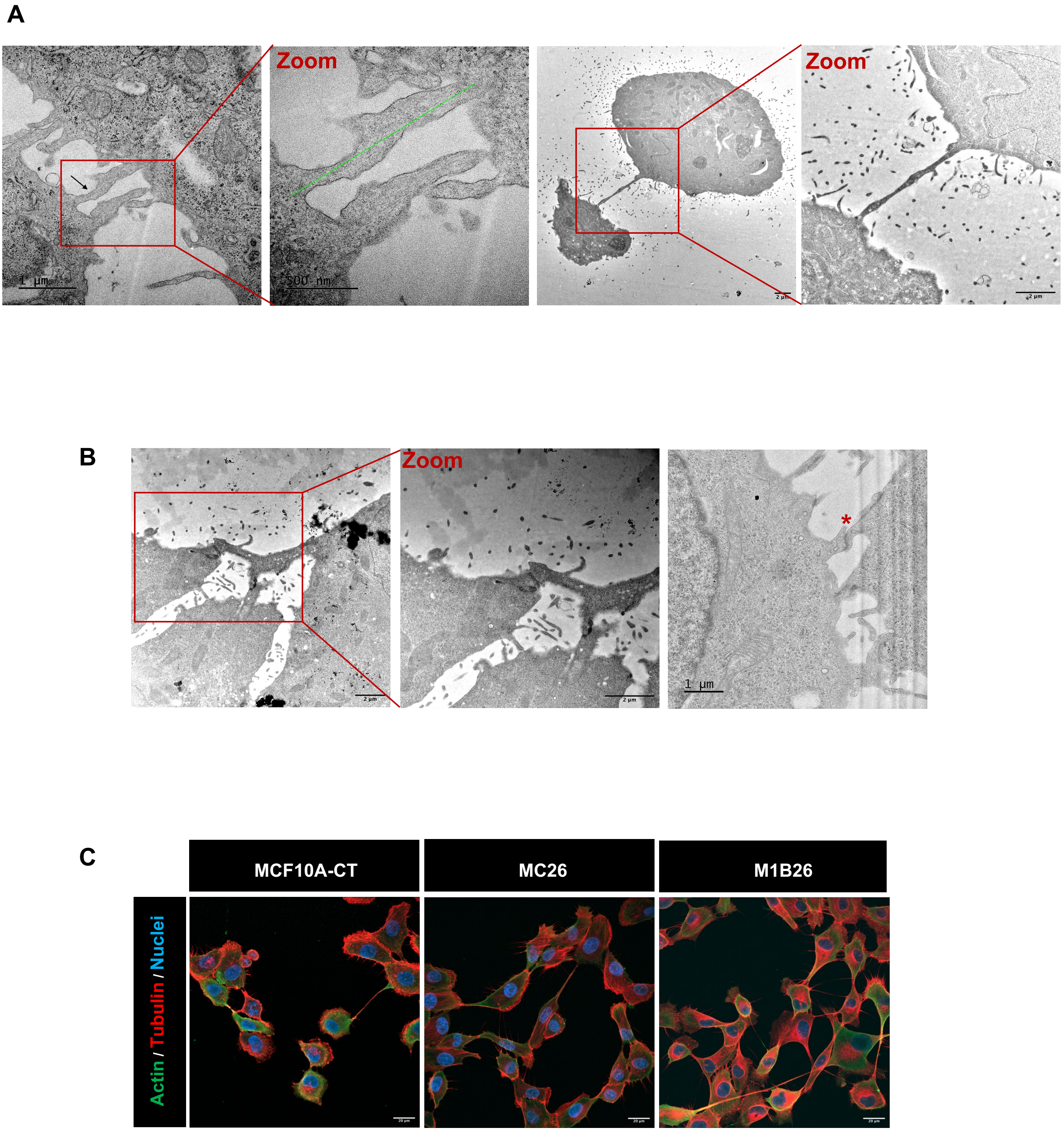


Supplementary Figure S2

**Supplementary Figure S2**. A, Transmission electron microscopy (TEM) images of open-ended TNTs linking two M1B26 cells. From left to right, scale bars : 1 µm, 500 nm, 2 µm, 2 µm. The green line indicates cytoplasmic continuum. B, TEM images of close-ended TNTs linking two M1B26 cells *via* a Gap junction. From left to right, scale bars, 2 µm, 2 µm, 1 µm. Red asterisk, Gap junction. C, Immunofluorescence images of actin (green) and tubulin (red) in indicated cell lines. Nuclei, blue. Scale bars, 20 µm.

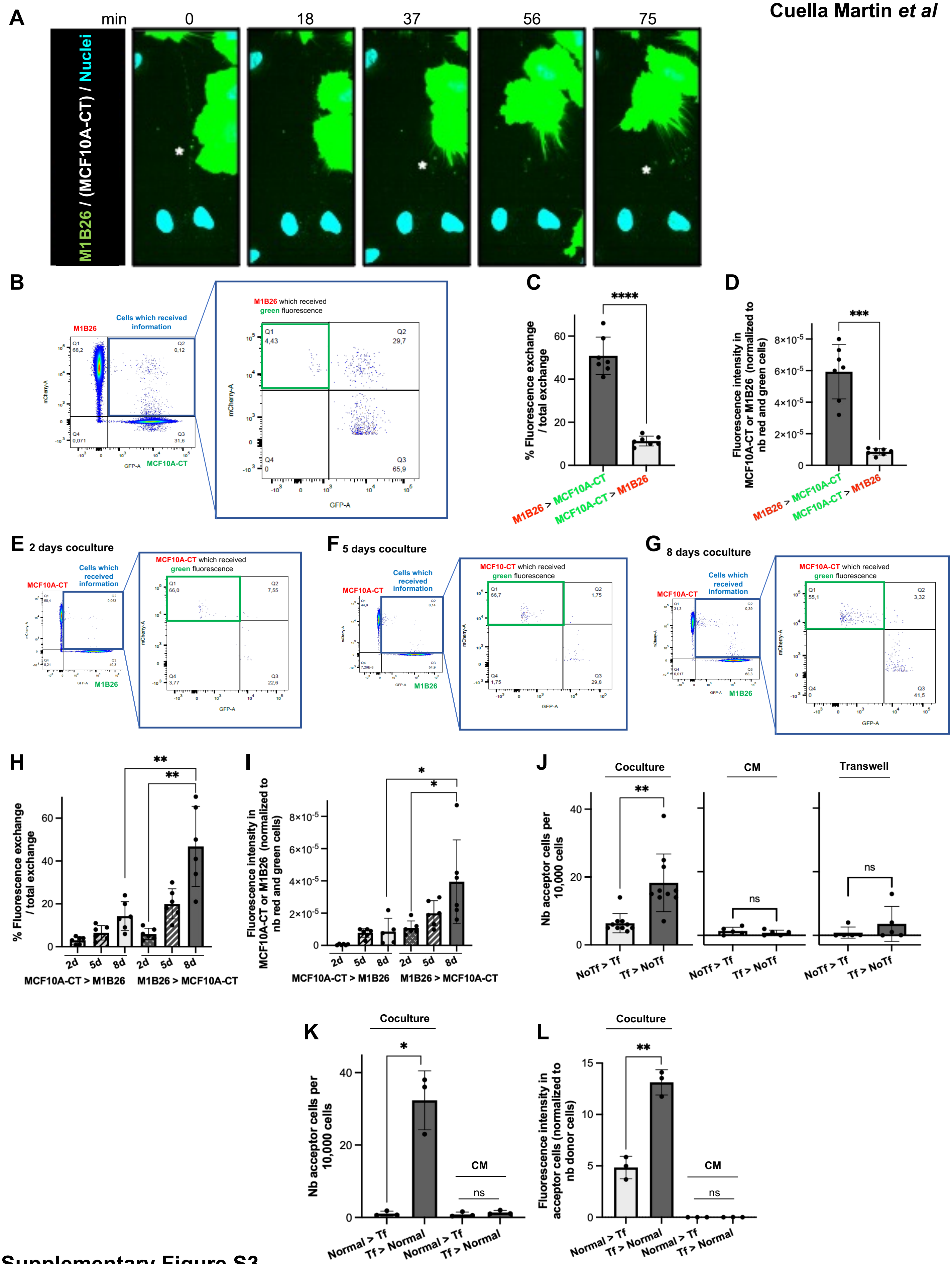


Supplementary Figure S3

**Supplementary Figure S3**. A, Time-lapse snapshots of green material trafficking within a TNT bulge (*), from GFP (green) M1B26 to MCF10A-CT cells after 48h in coculture. Nuclei, turquoise blue. Scale bar, 100 µm. Images were recorded for 75 min, with one image every 18 min. For movie source data, see Movie S4. B, Flow cytometry analysis of an 8-day coculture of red M1B26 and green MCF10A-CT cells. C,D, Quantification of the percentage of exchange (C) and fluorescence intensity (D), of the indicated transfer. Error bars depict SD. Statistical significance was calculated using an unpaired Welch's two-sided t test ($n = 7$ biological replicates; *** $p < 0.001$; **** $p < 0.0001$). E,F,G, Flow cytometry analysis of 2 (E), 5 (F) or 8 (G) days of coculture of green M1B26 and red MCF10A-CT cells. H,I, Percentage of material exchange (H) and fluorescence intensity (I), of the indicated transfer after 2, 5 or 8 days of coculture. Error bars are SD. Statistical significance was calculated using a one-way ANOVA with post-hoc Tukey Honest Significance Difference test ($n = 6$ biological replicates; * $p < 0.05$; ** $p < 0.01$). For clarity, only statistical significance is shown. J, Quantification of the number of acceptor cells *per* 10,000 cells, in standard (10 biological replicates) or Transwell cocultures (5 biological replicates), or in conditioned medium (5 biological replicates), for the indicated transfer involving MCF10A-CT (NoTf) and M1B26 (Tf) cells. Error bars are SD. Statistical significance was calculated using an unpaired Welch's two-sided t test (ns, not significant; ** $p < 0.01$). K,L, Quantification of the number of acceptor cells *per* 10,000 cells (K), and fluorescence intensity (L), in cocultures or with conditioned medium, for the indicated transfer involving normal human primary mammary (normal) and M1B26 cells (Tf). Error bars are SD. Statistical significance was calculated using an unpaired Welch's two-sided t test ($n = 3$ biological replicates; ns, not significant; * $p < 0.05$; ** $p < 0.01$).

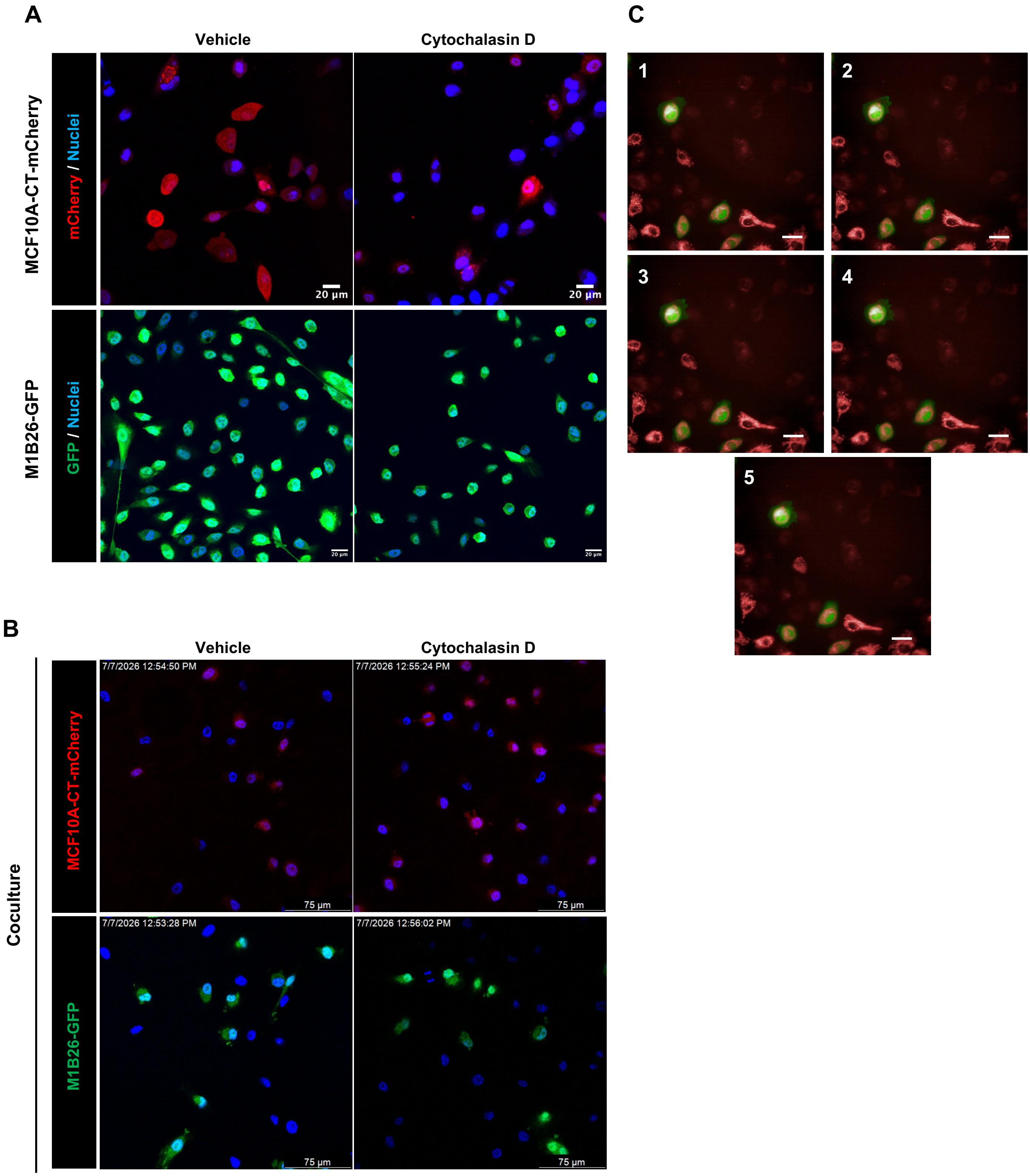


Supplementary Figure S4

**Supplementary Figure S4**. A, Fluorescence images of mCherry-labeled MCF10A-CT or GFP-labeled-M1B26 cells. Before fixation, cells were treated for 60 min with 1 µM cytochalasin D, or with DMSO (vehicle). Observations were performed by confocal microscopy in Airyscan mode, with an 40x oil objective. Scale bars, 20 µm. B, Fluorescence images of GFP-labeled-M1B26 and mCherry-labeled MCF10A-CT cells, after 48h in coculture. Before fixation, cells were treated for 60 min with 1 µM cytochalasin D, or with DMSO (vehicle). Scale bars, 75 µm. Nuclei, Hoechst labeling (blue). C, Snapshots from Movie S5 (images from 1 to 5). GFP-labeled M1B26 and unlabeled MCF10A-CT cells, cocultured for 48h, were labeled with Mitotracker™ DeepRed, before their treatment with 1 µM cytochalasin D for 60 min. Live imaging was performed on a confocal Revvity Opera Phenix time-lapse microscope with 40x water objective. Scale bar, 20 µm.

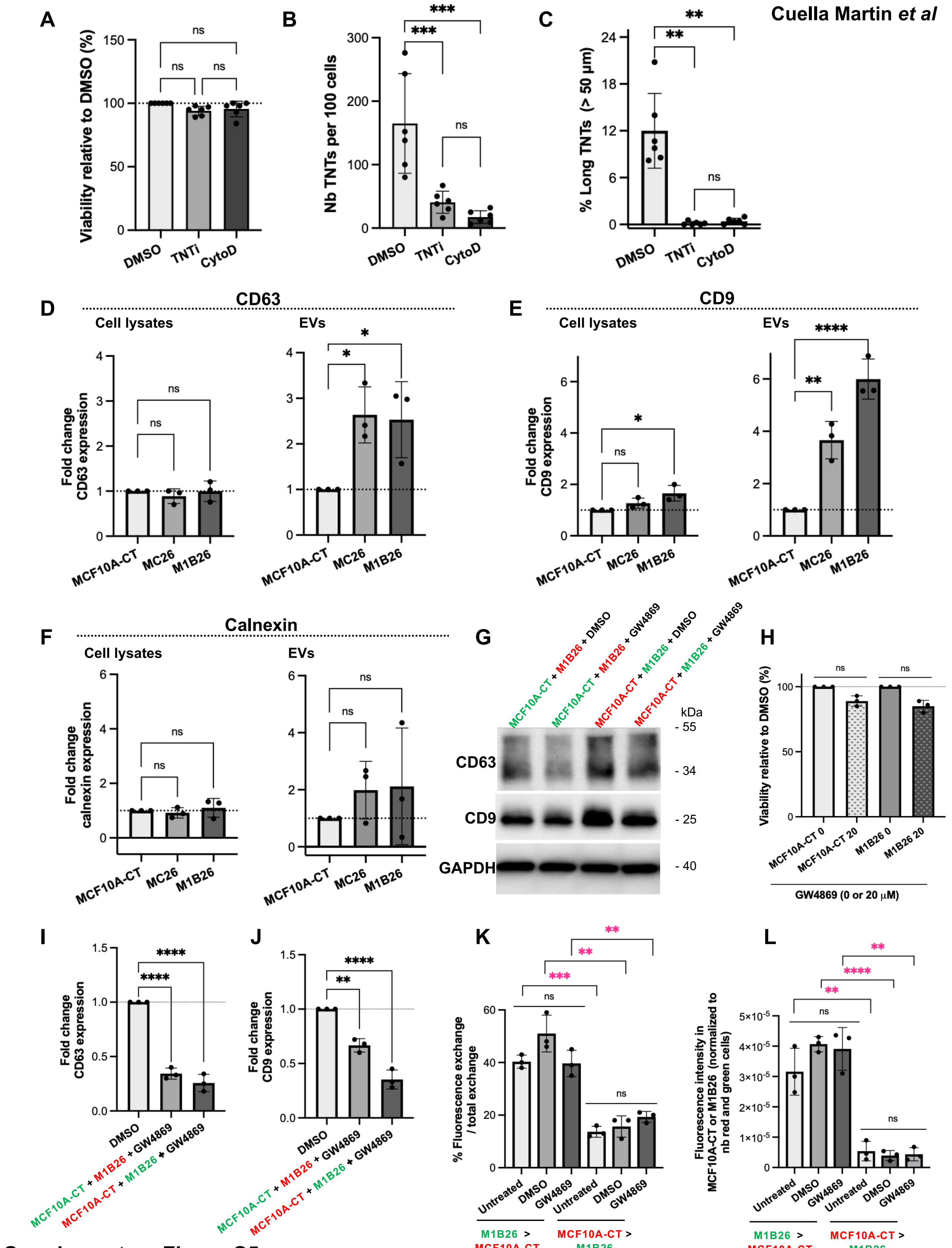


Supplementary Figure S5

**Supplementary Figure S5**. A, Histograms showing the viability of M1B26 cells treated for 1h with 20 μM TNTi or 1 μM cytochalasin D, relative to DMSO-treated cells, as measured with MTT assay (n = 6 biological replicates). B-C, Quantification of TNT number (B) and of % of long TNTs (> 50 μm, C) for M1B26 cells and indicated conditions; n = 6 biological replicates. Ten Z-stacks of 10 sections each were taken per condition, and 100 cells minimum *per* experiment were counted. TNTs were counted and measured in each Z-stack. Error bars depict SD. P-values were calculated with one-way ANOVA with post-hoc Tukey's Honest Significance Difference test (ns, not significant; ** $p < 0.01$; *** $p < 0.001$). D-F, Quantification of immunoblots of indicated proteins, expressed as fold change protein expression to MCF10A-CT cells set at 1. EVs, extracellular vesicles obtained by pelleting cell supernatants (see Methods). G, Representative immunoblot of CD63 and CD9 from EVs of indicated cellular conditions. GW4869, inhibitor of EVs biogenesis (20 μM). H, Histograms showing the viability of indicated cells, treated with 20 μM GW4869 for 24h, relative to DMSO (0); ns, not significant (n = 3 biological replicates). I-J, Quantification of immunoblots of CD63 (I) and CD9 (J), for indicated cellular conditions, with DMSO control set at 1. P-values were calculated with one-way ANOVA with post-hoc Tukey's Honest Significance Difference test (** $p < 0.01$; **** $p < 0.0001$). K-L, Quantification of the percentage of exchange (K; n = 3 biological replicates) and fluorescence intensity (L; n = 3 biological replicates), for indicated transfers. Error bars depict SD. Statistical significance was calculated using an unpaired Student's two-sided Welch's t test (ns, not significant; ** $p < 0.01$; *** $p < 0.001$).

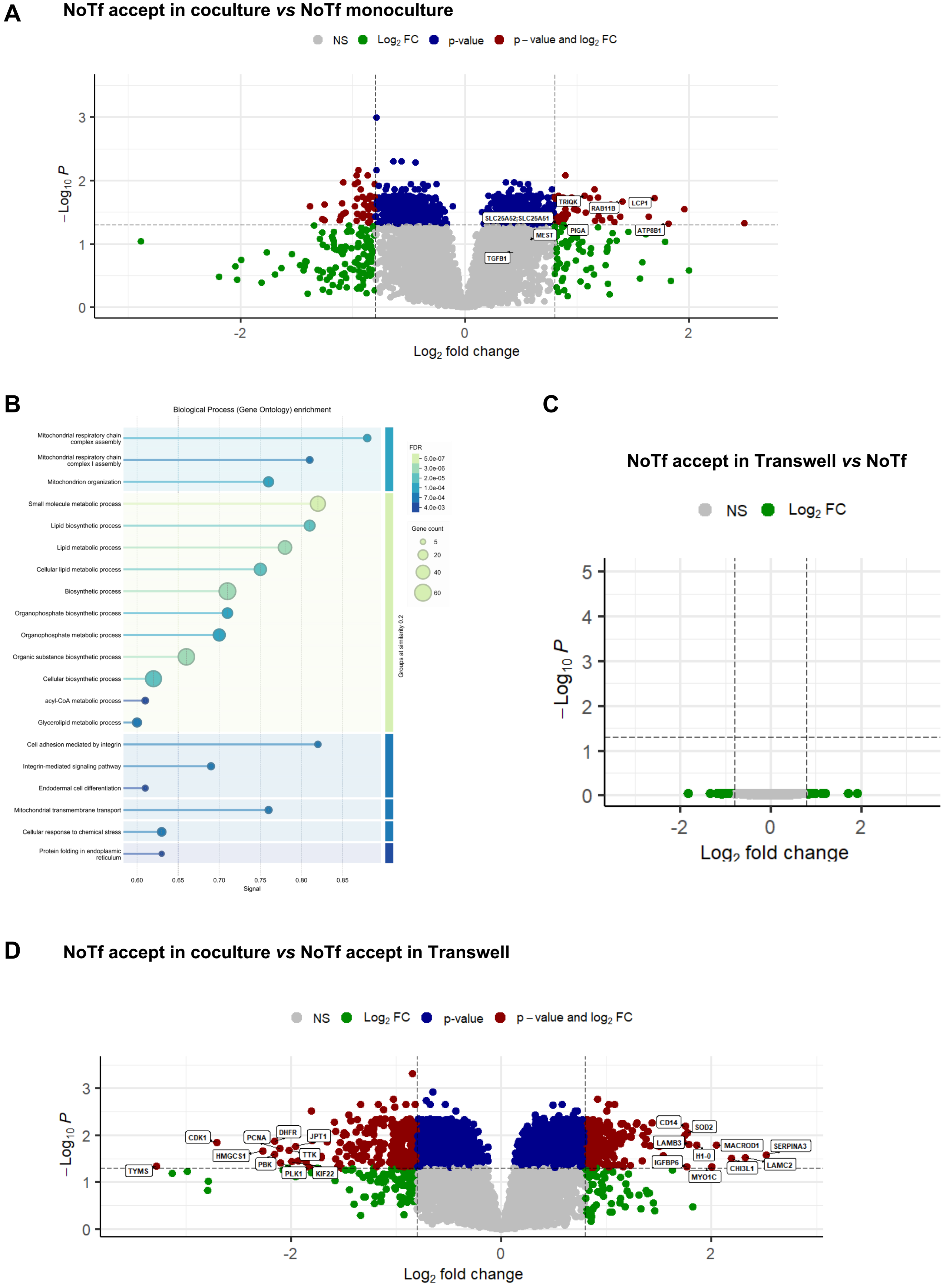


**Supplementary Figure S6**

**Supplementary Figure S6**. A, Volcano plot showing $Log_2$ enrichment of individual proteins in non-transformed acceptor cells (NoTf accept) in direct coculture, compared to non transformed (NoTf) cells in monoculture. The threshold for the adjusted p-value (with Benjamini-Hochberg procedure) was set at 0.05 and 0.8 for Log2 fold change. The Log2 fold change threshold was lowered to 0.5 in order to obtain a larger pool of hits, with a minimum of differences between groups. B, From a list of 193 proteins upregulated in NoTf accept cells (in coculture) compared to NoTf cells (in monoculture), STRING (https://string-db.org; RRID:SCR_005223) was used to generate the gene ontology analysis with the biological process subset. C,D, Volcano plots showing $Log_2$ enrichment of individual proteins in NoTf accept in Transwell coculture compared to NoTf cells in monoculture (C), and in NoTf accept cells in direct coculture compared to NoTf accept cells in Transwell coculture (D). The threshold for the adjusted p-value (with Benjamini-Hochberg procedure) was set at 0.05 and 0.8 for Log2 fold change. The Log2 fold change threshold was lowered to 0.5 in order to obtain a larger pool of hits, with a minimum of differences between groups.

Cuella Martin *et al*

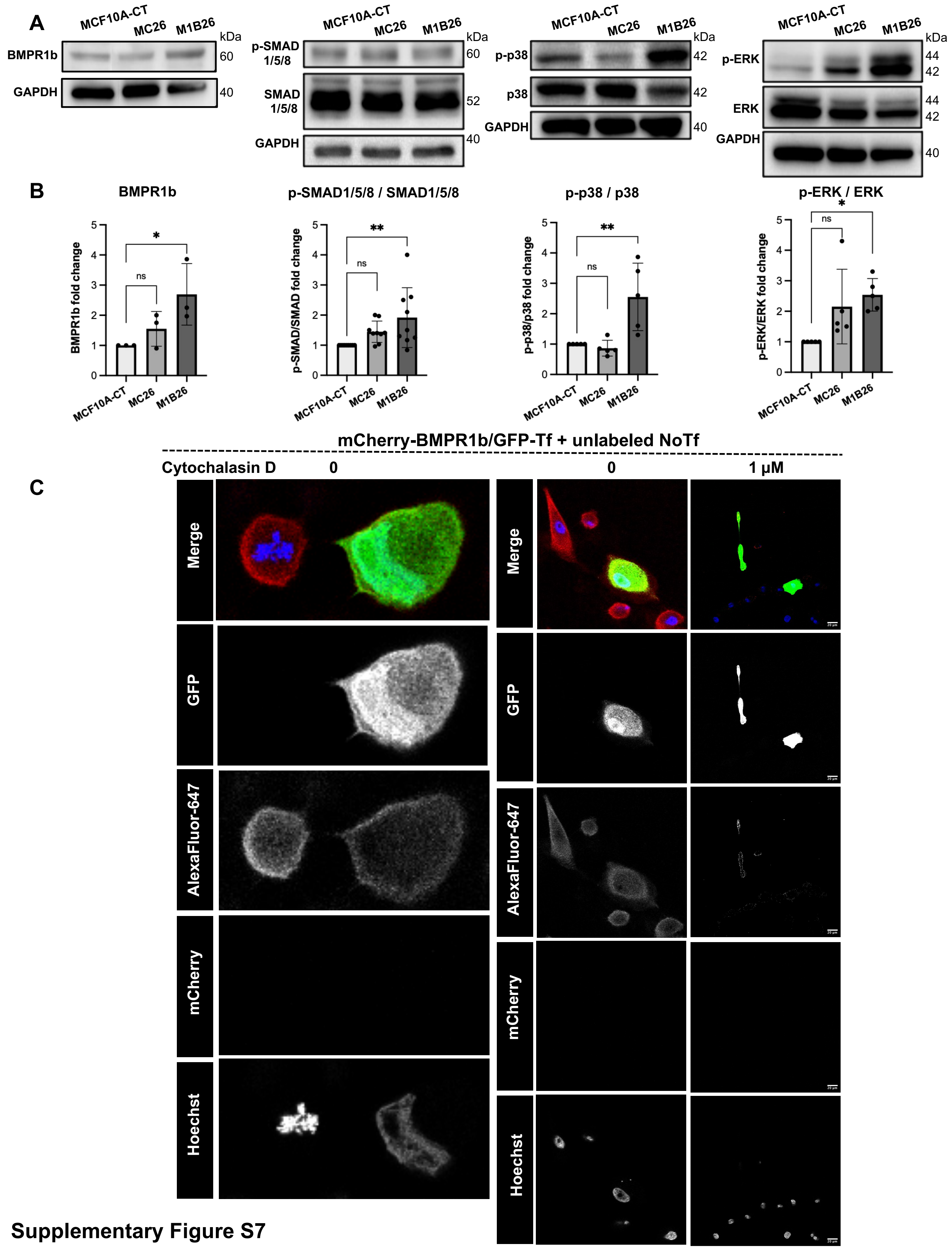


Supplementary Figure S7

**Supplementary Figure S7.** A, From left to right : Immunoblots of indicated proteins, in MCF10A-CT, MC26 and M1B26 cells. B, From left to right : quantification of immunoblots shown in A. Error bars are SD. Statistical significance was calculated using a one-way ANOVA with post-hoc Tukey's Honest Significance Difference test (biological replicates : n = 3 for BMPR1b, n = 9 for p-SMAD1/5/8/SMAD1/5/8, n = 5 for p-p38/p38 and p-ERK/ERK ; ns, not significant, * $p < 0.05$ ; ** $p < 0.01$). Data are mean ± SD. C, Representative immunofluorescence images of GFP-labeled M1B26 (Tf) cells (green) transfected with an mCherry-expressing BMPR1b construct, before their coculture for 48h with unlabeled MCF10A-CT (NoTf). GFP, green; antibody to mCherry with secondary antibody coupled to AlexaFluor® 647, red (see Methods); nuclei, blue. Right panels, cocultures treated for 1h with 1 µM cytochalasin D before fixation. All fluorescence channels were split for better visualization. Scale bar, 20 µm.

Cuella Martin *et al*

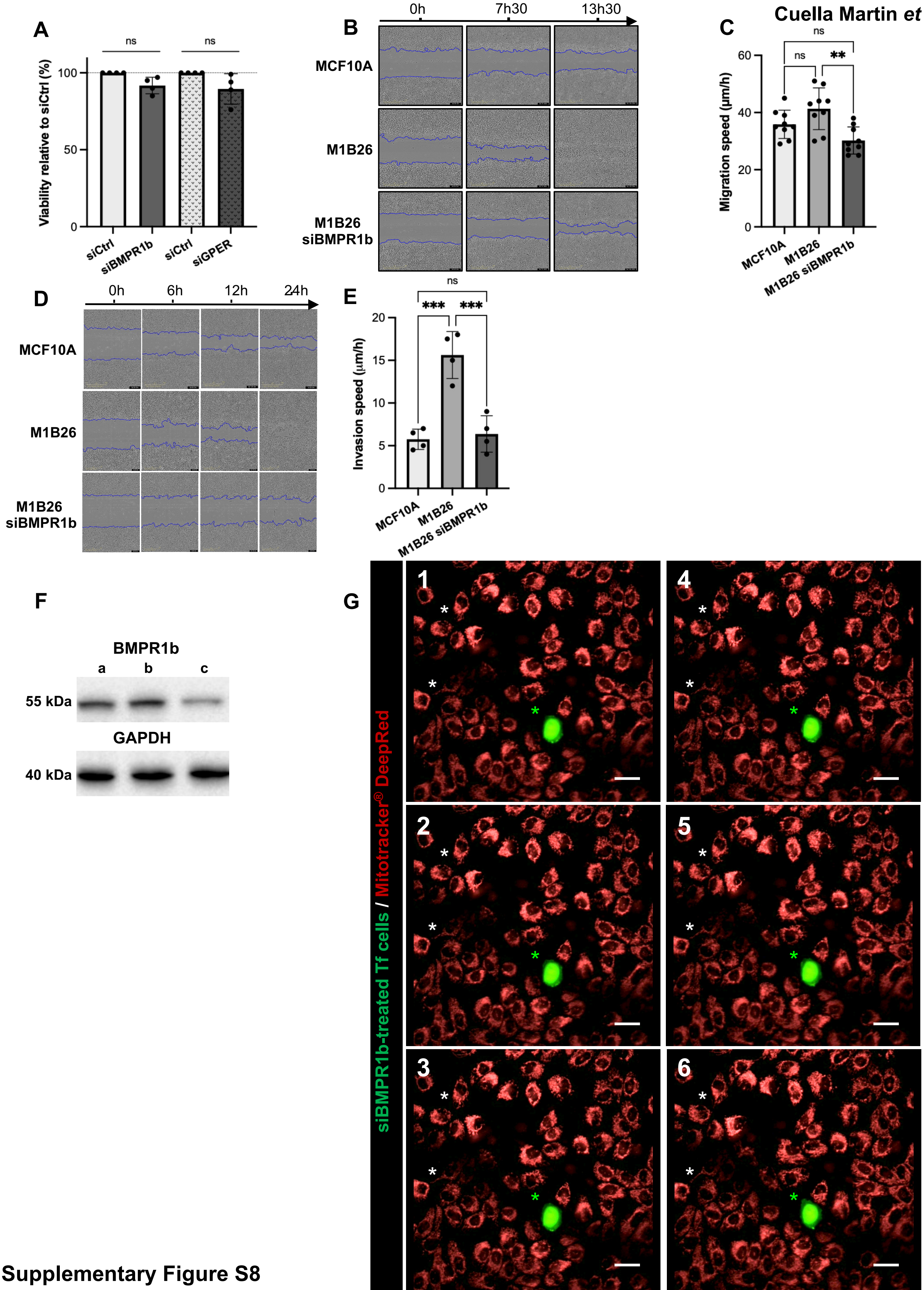


Supplementary Figure S8

**Supplementary Figure S8.** A, Histograms showing the viability of M1B26 transformed cells treated with control siRNA (siCtrl), siRNA targeting BMPR1b (siBMPR1b) for 48h or GPER (siGPER) for 72h, as measured with MTT assay (n = 4 biological replicates; ns = not significant). B, Observation of MCF10A, M1B26 and siBMPR1b-treated M1B26 migration for 13h30, after performing a scratch with a wound healing plate and live imaging during 24h by Incucyte® microscopy (representative images). C, Quantification of migration speed (µm/h) in MCF10A, M1B26, and siBMPR1b-treated M1B26. Data are mean±SD. Statistical significance was calculated using a one-way ANOVA with post-hoc Tukey's Honest Significance Difference test (n= 9 biological replicates; ** $p < 0.01$). D, Observation of MCF10A, M1B26 and siBMPR1b-treated M1B26 invasion for 24h, after performing a scratch with a wound healing plate and live imaging by Incucyte® microscopy (representative images). E, Quantification of invasion speed (µm/h) in MCF10A, M1B26 and siBMPR1b-treated M1B26. Data represent mean±SD. Statistical significance was calculated using a one-way ANOVA with post-hoc Tukey's Honest Significance Difference test (n = 4 biological replicates; ** $p < 0.01$, *** $p < 0.001$). F, Representative immunoblots of indicated proteins in Tf cells : a, untreated cells; b, cells treated with control siRNA; c, cells treated with siRNA targeting BMPR1b. G, Immunofluorescence of GFP-labeled Tf cells (green), treated for 36h with an siRNA targeting BMPR1b before their coculture with unlabeled NoTf cells. Mitochondria were stained in the coculture with Mitotracker® DeepRed (647 nm, red). The contrast of images was purposedly enhanced for better visualization. Numbers 1 to 6, successive snapshots extracted from Movie S6. White asterisks, mitochondria transiting in TNT-like structures between NoTf cells; green asterisks, GFP-labeled Tf cell, silenced for BMPR1b before coculture with NoTf cells, in close vicinity with NoTf cells.

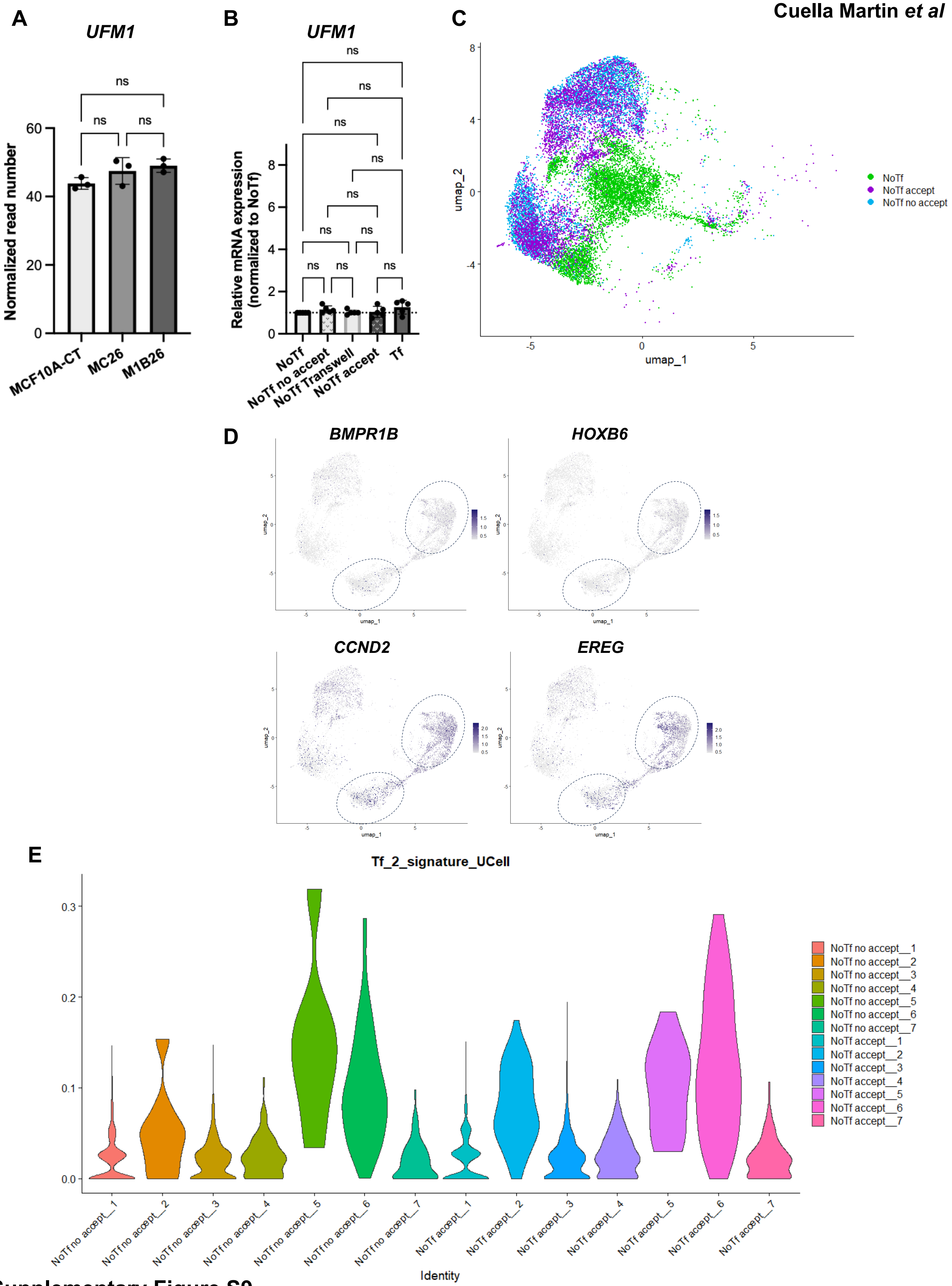


**Supplementary Figure S9**

**Supplementary Figure S9.** A, RNA-sequencing results showing the transcriptomic expression levels of UFM1 in indicated cell lines (n = 3 biological replicates). Data deposited on the Gene Expression Omnibus repository (RRID:SCR_005012), GSE186734 [6]. Error bars represent SD. Statistical significance was calculated using a one-way ANOVA with post-hoc Tukey's Honest Significance Difference test (ns, not significant). B, RT-qPCR measurements of UFM1 mRNA in indicated cell populations, with the expression in NoTf cells set at to 1. Error bars represent SD. Statistical significance was calculated using a one-way ANOVA with post-hoc Tukey's Honest Significance Difference test (n = 5 biological replicates; ns, not significant). C, U-maps comparing single cell transcriptomic profiles of NoTf, NoTf no accept and NoTf accept cells. D, Expression profiles at single cell resolution of *BMPR1B, HOXB6, CCND2* and *EREG* in Tf cells. E, Transcriptomic profile heterogeneity across MCF10A models and post TNT-mediated material transfer. UCell signature scoring plots. Tf_2_signature_UCell corresponds to the 30 most highly expressed genes identified in cluster 2 of Tf cells. The plots show the distribution of this signature across clusters of NoTf cells, distinguishing cells that have captured material from Tf cells (NoTf accept) from those that have not (NoTf no accept). Data were deposited on the Gene Expression Omnibus repository (RRID:SCR_005012), GSE343015.

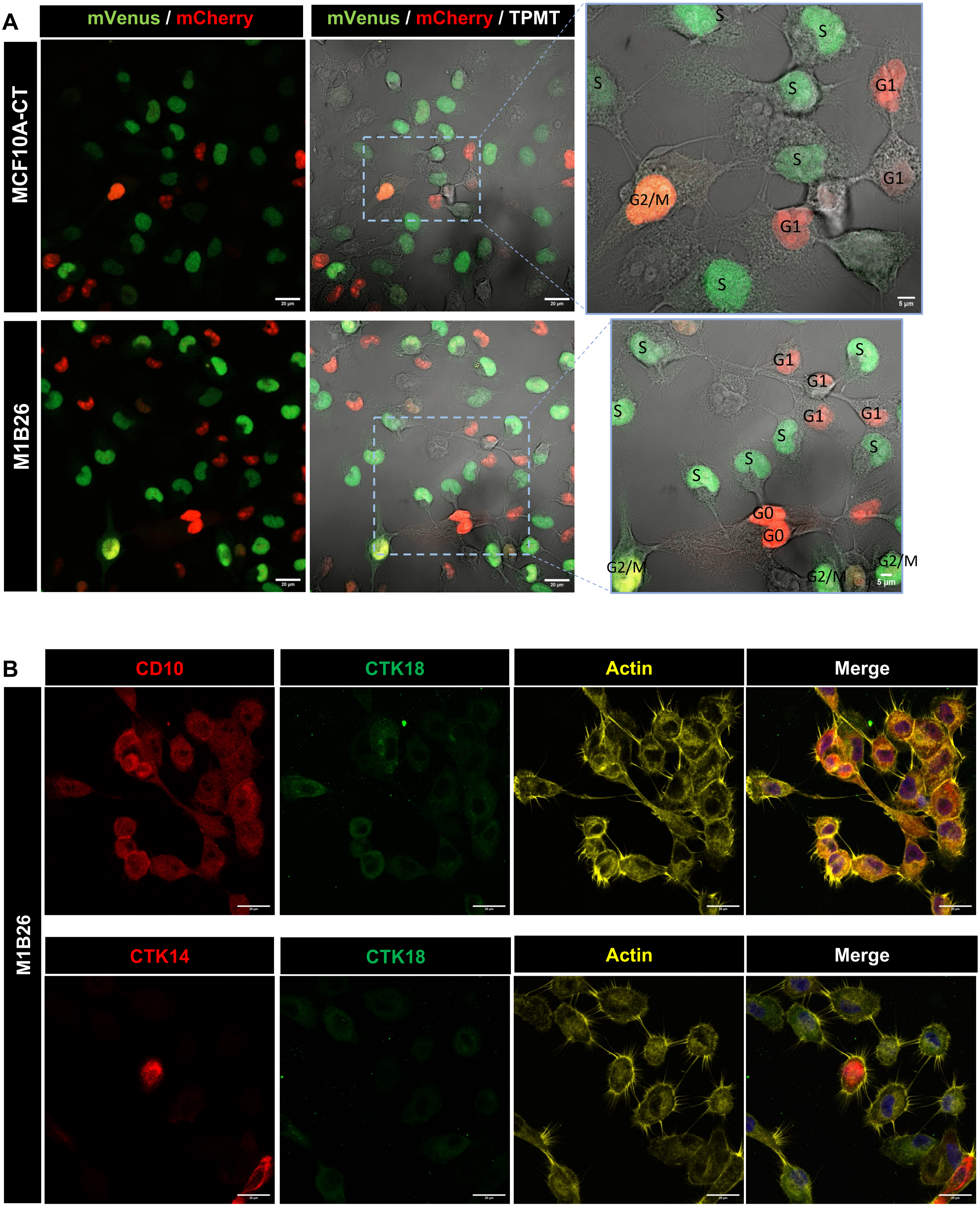


Supplementary Figure S10

**Supplementary Figure S10.** A, Representative images of MCF10A-Fucci and M1B26-Fucci [32] cells by Airsycan confocal microscopy with transmitted light (TPMT). *mCherry+++: G0 phase; mCherry+: G1 phase; mVenus+/mCherry+: G2/M phase, mVenus+: S phase; Non-fluorescent: G1-S transition. Scale bar, 20 µm, or 5 µm in zooms. B, Representative images of stem (CD10, red, top), myoepithelial (CTK14 or cytokeratin 14, red, bottom) and epithelial (CTK18 or cytokeratin 18, green) markers in MCF10A and M1B26 cells, immunostained for actin (yellow) and connected by TNTs, obtained by Airyscan confocal microscopy. Nuclei, Hoechst (blue). Scale bar, 20 µm.

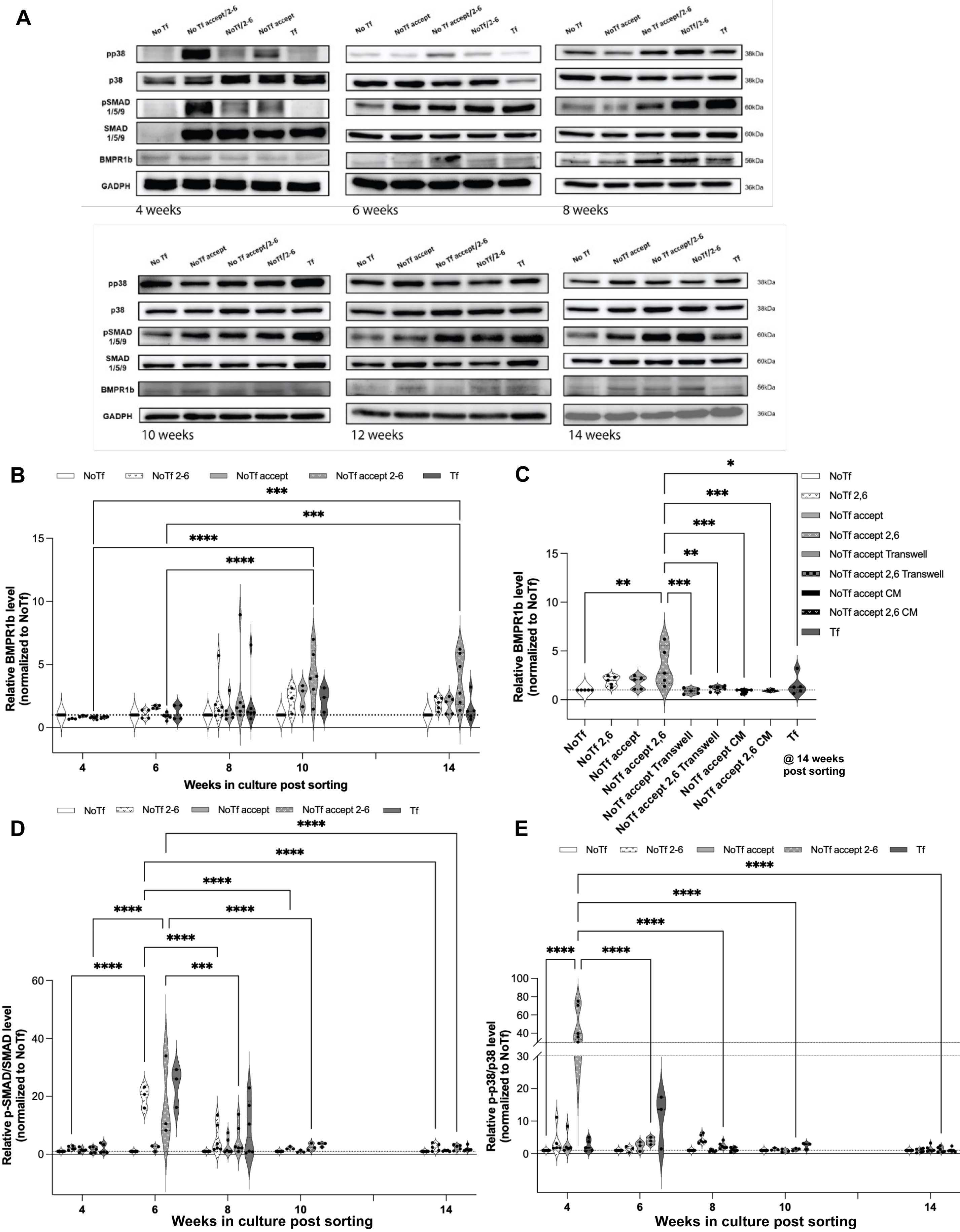


Supplementary Figure S11

**Supplementary Figure S11**. A, Representative immunoblots of indicated proteins in indicated cells, grown for indicated weeks in the absence or presence of BMP2/IL6 (2-6). B, Quantification of BMPR1b immunoblots, from lysates of cells grown for indicated time post flow cytometry sorting (see schematic Figure 8A). Violin plots of 3 to 6 biological replicates, with NoTf and NoTf 2-6, MCF10A-CT cells grown in the absence or presence of BMP2/IL6, respectively; NoTf accept and NoTf accept 2-6, MCF10A cells grown in coculture with M1B26 cells, and cultured in the absence or presence of BMP2/IL6 post-sorting, respectively; Tf, M1B26 cells. NoTf conditions are set at 1. Statistical significance was calculated using a two-way ANOVA with Tukey's Honest Significance Difference multiple comparison test. For clarity, only significant comparisons are shown.

C, Quantification of BMPR1b immunoblots performed on indicated cells, 14 weeks post sorting, with NoTf conditions set at 1. Violin plot of 5 biological replicates. Statistical significance was calculated using a one-way ANOVA with post-hoc Tukey's Honest Significance Difference test, with only significant comparisons shown (* $p < 0.05$; ** $p < 0.01$; *** $p < 0.001$). Legend as in B, with : Transwell, coculture in Transwell; CM, NoTf (MCF10A-CT) cells grown in conditioned medium of M1B26 cells, sorted and grown in the absence or presence of BMP2/IL6. D-E, Quantification of p-SMAD1/5/8/SMAD1/5/8 (D) and p-p38/p38 (E) immunoblots, on same samples as in A and at indicated weeks post sorting. Statistical analyses were performed as in B.

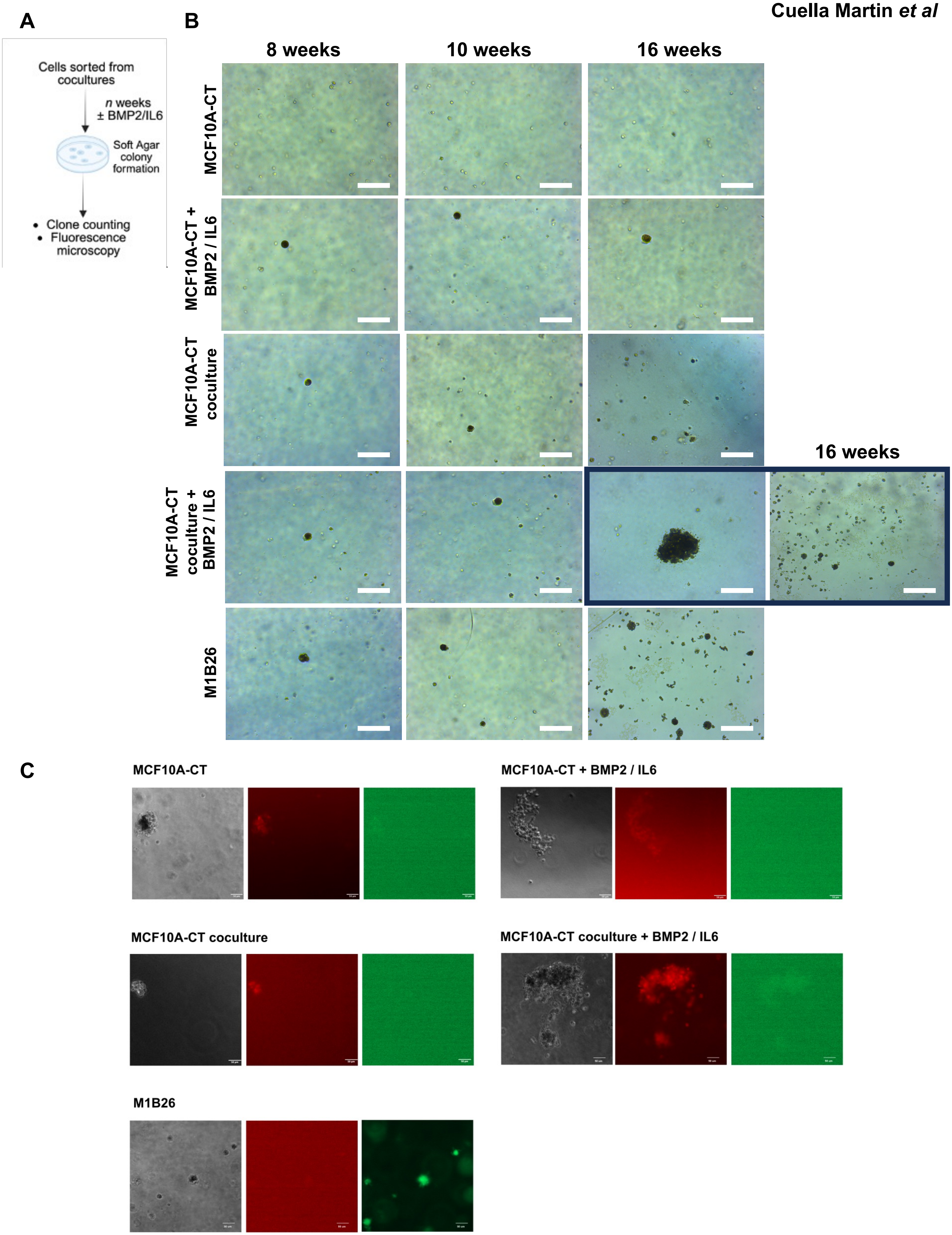


Supplementary Figure S12

**Supplementary Figure S12**. A, Schematic of long-term soft agar colony formation assays. B, Representative bright-field images of soft agar clones after indicated weeks in the absence or presence of BMP2/IL6. Pictures with a dark frame are representative of big or small clones observed for the indicated cell condition. Scale bar represents 200 μm. C, Representative images of soft agar clones obtained from indicated cells and conditions, and observed by epifluorescence microscopy. After sorting by flow cytometry, cells were grown in the absence or presence of BMP2/IL6 for several weeks (see Figure 8A), and at given time points, collected to perform soft agar colony formation assay (see panel A). Clones shown here were formed from cells grown for 16 weeks post sorting. From left to right : bright field, mCherry, GFP. Scale bars are 50 μm.

**Supplementary Table S1**. The ten most differentially expressed genes (from proteomic analyses) between NoTf accept and NoTf no accept cells, in direct coculture conditions, showing $Log_2$ enrichment of individual proteins (LogFC) and corresponding adjusted p-value with FDR (see legend to Supplementary Figure S6 for more details).

| UP | | | | | DOWN | | | |
|---|---|---|---|---|---|---|---|---|
| Proteins | LogFC | Adjusted p value | FDR | | Proteins | LogFC | Adjusted p value | FDR |
| ATP8B1 | 2.09279285090789 | 0.000411271663574841 | 0.0136694846866256 | | TMA7 | -0.967897673428647 | 0.00711927694228354 | 0.0425927067413436 |
| RAB11B | 1.3714054705064 | 0.000626507681921401 | 0.0157617717578257 | | PEBP1 | -0.992672308990052 | 0.000148184132312685 | 0.010254537822385 |
| LCP1 | 1.09840574413178 | 0.00154609002947331 | 0.0219662052490839 | | LGMN | -0.994280406934229 | 0.000616181168435651 | 0.0157617717578257 |
| MEST | 0.932759743282637 | 0.00188098627865132 | 0.023309540980721 | | ABHD14B | -1.00315026985703 | 0.000980307865197619 | 0.018125067074596 |
| TGFB1 | 0.865460115477054 | 0.00448296638926345 | 0.0342292515516622 | | PHPT1 | -1.01510146569795 | 0.00356732439665008 | 0.0307447685936627 |
| SLC25A52;SLC25A51 | 0.844728235272825 | 0.00690767760995108 | 0.0418102459418638 | | JPT2 | -1.10698538807139 | 0.00935740869041231 | 0.0487956300793808 |
| PIGA | 0.837875077610601 | 0.00443809956925508 | 0.0340571776719389 | | JPT1 | -1.10835843431927 | 0.00793600149391597 | 0.0448843931437067 |
| TRIQK | 0.827399246777439 | 0.00293132702439602 | 0.0280707500273152 | | TMSB10 | -1.14967105039942 | 0.00540682517524317 | 0.0369468666022315 |
| SLC22A4 | 0.776648784138638 | 0.00150878846847539 | 0.0217758203883326 | | ZNF706 | -1.15566233765069 | 0.00111603705555586 | 0.0190657244022509 |
| TMEM97 | 0.72583530827107 | 0.00976058972544308 | 0.0496690455478472 | | EEF1AKMT2 | -1.27111422803223 | 0.00946055818632448 | 0.0489747295783692 |

**Supplementary Table S2**. The ten most differentially expressed genes (from proteomic analyses) between NoTf accept in direct coculture and NoTf accept in Transwell coculture conditions, showing $Log_2$ enrichment of individual proteins (LogFC) and corresponding adjusted p-value with FDR (see legend to Supplementary Figure S6 for more details).

| UP | | | | | DOWN | | | |
|---|---|---|---|---|---|---|---|---|
| **Protein** | **LogFC** | **Adjusted p value** | **FDR** | | **Protein** | **LogFC** | **Adjusted p value** | **FDR** |
| SERPINA3 | 2.52280933849719 | 0.00535426160167644 | 0.0260595536346162 | | JPT1 | -1.95890497354932 | 0.0026510562963446 | 0.017271714851675 |
| LAMC2 | 2.3234530990449 | 0.00671123053131829 | 0.0297826525508751 | | KIF22 | -1.9960721026904 | 0.00971764170582591 | 0.0370559831125849 |
| CHI3L1 | 2.18947902717664 | 0.00727017238146173 | 0.0310809074549274 | | TTK | -2.02038496716218 | 0.00363301502290389 | 0.0211348472887102 |
| MACROD1 | 2.04828979371154 | 0.00240255771521197 | 0.0164486348091409 | | PLK1 | -2.09690800691763 | 0.0106792923363495 | 0.0394696070027224 |
| MYO1C | 2.0046600551578 | 0.0143021854781629 | 0.0475205948265113 | | PCNA | -2.10549674961135 | 0.00314193588565591 | 0.0193172329444116 |
| H1-0 | 1.85763513264292 | 0.00242570948335453 | 0.0165435708017873 | | PBK | -2.15384894310531 | 0.00508833653275616 | 0.0252538819421679 |
| LAMB3 | 1.78650743040783 | 0.00216132909781278 | 0.0156239730978036 | | DHFR | -2.15668666737884 | 0.0015530984577745 | 0.0135317025777003 |
| SOD2 | 1.77111253782108 | 0.000523439457102529 | 0.00883342599161415 | | HMGCS1 | -2.26815542277271 | 0.003911088063124 | 0.0217938425534674 |
| IGFBP6 | 1.76618122025904 | 0.0139229645066205 | 0.0468061169993795 | | CDK1 | -2.70314055151148 | 0.0017239046757659 | 0.0141924949054157 |
| CD14 | 1.75212297141304 | 0.000191011658635565 | 0.00639126803331301 | | TYMS | -3.27748800106074 | 0.0134778717596743 | 0.0457920533805404 |

**Supplementary Table S3**. Pathway analyses were performed using STRING (https://string-db.org/; RRID:SCR_005223), with the Biological Process category from the Gene Ontology collection. For the search of BMP and p38 pathways targets, a combination of several pathways from different collections have been compiled (GOBP_POSITIVE_REGULATION_OF_BMP_SIGNALING_PATHWAY ; GOBP_REGULATION_OF_BMP_SIGNALING_PATHWAY ; GOBP_RESPONSE_TO_BMP ; PID_BMP_PATHWAY ; BIOCARTA_P38MAPK_PATHWAY ; GOBP_P38MAPK_CASCADE ; GOBP_REGULATION_OF_P38MAPK_CASCADE ; REACTOME_P38MAPK_EVENTS ; WP_P38_MAPK_SIGNALING).

| #Category | Term ID | Term description | Observed gene counts | Background gene counts | Strength | Signal | False discovery rate (FDR) | Matching proteins in network (IDs) | Matching proteins in network (labels) |
|---|---|---|---|---|---|---|---|---|---|
| GO Process | GO:0007166 | Cell surface receptor signaling pathway | 7 | 2040 | 0.93 | 0.67 | 0.0068 | 9606.ENSP00000264110,9606.ENSP00000280377,9606.ENSP00000339191,9606.ENSP00000348461,9606.ENSP00000356070,9606.ENSP00000379975,9606.ENSP00000414147 | ATF2,USP15,CAV1,RAC1,MAPKAPK2,ILK,SPART |
| GO Process | GO:0071363 | Cellular response to growth factor stimulus | 5 | 473 | 1.42 | 1.01 | 0.0068 | 9606.ENSP00000264110,9606.ENSP00000280377,9606.ENSP00000339191,9606.ENSP00000356070,9606.ENSP00000414147 | ATF2,USP15,CAV1,MAPKAPK2,SPART |
| GO Process | GO:0007167 | Enzyme-linked receptor protein signaling pathway | 5 | 641 | 1.28 | 0.92 | 0.0075 | 9606.ENSP00000264110,9606.ENSP00000280377,9606.ENSP00000348461,9606.ENSP00000356070,9606.ENSP00000414147 | ATF2,USP15,RAC1,MAPKAPK2,SPART |
| GO Process | GO:0006468 | Protein phosphorylation | 5 | 736 | 1.22 | 0.82 | 0.0119 | 9606.ENSP00000264110,9606.ENSP00000280377,9606.ENSP00000339191,9606.ENSP00000356070,9606.ENSP00000379975 | ATF2,USP15,CAV1,MAPKAPK2,ILK |
| GO Process | GO:0030509 | BMP signaling pathway | 3 | 92 | 1.9 | 1.03 | 0.0156 | 9606.ENSP00000264110,9606.ENSP00000280377,9606.ENSP00000414147 | ATF2,USP15,SPART |
| GO Process | GO:0030510 | Regulation of BMP signaling pathway | 3 | 104 | 1.85 | 0.97 | 0.0191 | 9606.ENSP00000339191,9606.ENSP00000379975,9606.ENSP00000414147 | CAV1,ILK,SPART |
| GO Process | GO:0038066 | p38MAPK cascade | 2 | 13 | 2.58 | 1.01 | 0.0223 | 9606.ENSP00000264110,9606.ENSP00000356070 | ATF2,MAPKAPK2 |
| GO Process | GO:0071310 | Cellular response to organic substance | 6 | 2019 | 0.86 | 0.51 | 0.0356 | 9606.ENSP00000264110,9606.ENSP00000280377,9606.ENSP00000339191,9606.ENSP00000356070,9606.ENSP00000379975,9606.ENSP00000414147 | ATF2,USP15,CAV1,MAPKAPK2,ILK,SPART |

**Supplementary Table S4**. Primers for RT-qPCR used in this study.

| | |
|---|---|
| *HPRT* FW | TGACCTTGATTTATTTTGCATACC |
| *HPRT* RV | CGAGCAAGACGTTCAGTCCT |
| *TBP* FW | TGTATCCACAGTGAATCTTGGTTG |
| *TBP* RV | GGTTCGTGGCTCTCTTATCCTC |
| *BMPR1B* FW | CTGTGGTCACTTCTGGTTGC |
| *BMPR1B* RV | TTCCTTTCTGTGCAGCATTC |
| *EREG* FW | TCTGCCTGGGTTTCCATCTT |
| *EREG* RV | ATTGACACTTGAGCCACACG |
| *HOXB6* FW | CGTGCAACAGTTCCTCCTTT |
| *HOXB6* RV | GCGTCAGGTAGCGATTGTAGT |
| *CCND2* FW | TGCTGGAGTGGGAACTGGTG |
| *CCND2* RV | ACACAGAGCAATGAAGGTCT |
| *UFM1* FW | AAAGTTCCTGCTGCAACAAGTGC |
| *UFM1* RV | ACACGATCTCTAGGAATAATCCGC |
| *HRAS* FW | ACGCACTGTGGAATCTCGGCAG |
| *HRAS* RV | TCACGCACCAACGTGTAGAAGG |
